# Conditions enabling the persistence of cooperating ribozymes without cellular encapsulation

**DOI:** 10.1101/2025.06.10.658808

**Authors:** Zhen Peng, Alex M. Plum, Rahul Kartha, Emily M. Jacobson, David A. Baum

**Affiliations:** Wisconsin Institute for Discovery, University of Wisconsin–Madison, Madison, WI 53706, USA; Department of Engineering Physics, University of Wisconsin–Madison, Madison, WI 53706, USA; Department of Computer Sciences, University of Wisconsin–Madison, Madison, WI 53706, USA; Department of Botany, University of Wisconsin–Madison, Madison, WI 53706, USA

## Abstract

Explaining the origin of molecular systems composed of cooperating polymer sets that confer both metabolic and information processing functions is a key challenge in origins-of-life research. A related puzzle is the emergence of polymers of sufficient length to confer complex functions such as the RNA-dependent RNA polymerization with proofreading. Addressing these issues using computational models of template-guided replicating polymer systems is generally constrained by the exponential increase in diversity as the length of polymers increases. In this study, inspired by the computer game Tetris® and the **P**olymerase **C**hain **R**eaction (PCR) technique, we developed an abstract computational model of cooperative replicating polymer systems that avoids tracking all potential sequences. Using this model, we explored cooperative chemical ecosystems consisting of catalytic polymers conferring functions analogous to kinases, ligases, and mutation inhibitors. We show that prebiotic environments with micro-compartments with local exchanges enable multilevel selection that facilitates the survival of cooperating polymers. The ability of cooperative systems to persist is sensitive to intrinsic properties of catalysts such as catalytic efficiency and extrinsic factors such as dilution rate. These results provide a roadmap for future studies that look not just at persistence but also at the stepwise, *de novo* emergence of chemical ecosystems with both metabolic and information-processing capabilities.

## 1. Introduction

In life as we know it, multiple cooperating biochemical functions, each conferred by distinct catalytic polymers, are integrated into a reaction system capable of collective self-replication and adaptive evolution. A key challenge in origins-of-life research is revealing how such multi-functional cooperative systems can originate in plausible prebiotic environments.

Although living organisms usually use proteins to catalyze reactions and nucleic acids to encode genetic information, it has long been recognized that some RNAs (i.e., ribozymes) are capable of catalyzing reactions, resulting in the RNA-World hypothesis that there was a stage in abiogenesis when RNA polymers both acted as templates and undertook most or all necessary catalytic functions for collective self-replication (Cech, 2012; Gilbert, 1986; Nielsen et al., 1991; Robertson and Joyce, 2012; Schöning et al., 2000). A first necessary function is catalyzing the synthesis of activated nucleotides from simpler and more abundant environmental resources, since accumulating a diverse and concentrated supply of activated nucleotides is otherwise unlikely in plausible prebiotic environments (Bregestovski, 2015; Dagar et al., 2020). This function may be seen as a basic form of metabolism. A second necessary function is catalyzing templated-mediated replication of RNA strands by consuming activated nucleotides, which requires that replication be accurate enough to regenerate functional ribozymes at a higher rate than they are lost due to degradation or dilution. This function may be seen as a primitive form of genetics.

Biochemists have used *in vitro* selection to generate ribozymes capable of replicating DNAs or RNAs through polymerization or ligation (Breaker, 1997; Ekland and Bartel, 1996; Martin et al., 2015; Silverman, 2008; Zaher and Unrau, 2007). As a result, many origins-of-life researchers assume the primordial existence of a single “self-replicating” ribozyme, namely one that can catalyze the copying of its reverse-complement into a copy of itself (Cech, 1986). However, it appears that ribozymes with accurate RNA-dependent RNA polymerase activity would likely need to be hundreds or even thousands of nucleotides long, but such long ribozymes would be unlikely to emerge alongside their reverse-complements without prior catalyzed polymerization (Eigen, 1971; Eigen and Schuster, 1978a, 1978b, 1977; Higgs and Lehman, 2015). To resolve this paradox, theories have been developed that, instead of relying on a single long ribozyme, imagine the co-occurrence of multiple short catalytic polymers that cooperate in such a way as to equip the entire polymer system with the capacity for efficient self-replication. The best-known such models are the hypercycle theory (Eigen and Schuster, 1978a, 1978b, 1977) and autocatalytic set theory (Hordijk and Steel, 2016, 2013; Kauffman, 1986). Both have proven challenging to simulate computationally because standard approaches require that the reaction rates (for mass action kinetics) or propensities (for stochastic models like Gillespie algorithm (Gillespie, 2007, 1977, 1976)) of all possible reactions are tracked and updated as quantities of reactants and catalysts change. Given that the number of possible reactions is determined by the diversity of polymer sequences allowed, which grows exponentially with maximum polymer length, traditional simulation approaches face a severe challenge. Therefore, to simulate the possible stepwise origins of genetic life from the ground-up, it is necessary to develop a new method that circumvents the immense combinatorial space of polymer sequences.

In this study, we developed a discrete stochastic model in a spatial context, as suggested by the stochastic corrector model (Grey et al., 1995; Szathmáry and Demeter, 1987; Zintzaras et al., 2002) and the theory of surface metabolism (Wächtershäuser, 1997, 1988). Inspired by the computer program Tetris®, we developed a strategy where the computational load is primarily set by reactor size and the count of realized catalytic polymer molecules in the system rather than the number of possible polymer sequences. Thus, our model circumvents the combinatorial explosion problem and can efficiently simulate stochastic dynamics relevant to the origins of cooperative polymer systems.

Our reaction system includes a highly simplified representation of both metabolism and genetics in a putative prebiotic environment and explores the stepwise emergence and complexification of cooperating ribozymes. We started with a basic model that uses two instead of the standard four bases and considered two primary catalytic functions: converting environmental resources into activated monomers (“metabolism”) and ligating activated monomers into polymers in a template-guided manner (“genetics”). In subsequent simulations, we extended this basic model with a “mutation inhibitor” function that serves to improve the fidelity of template-guided ligation and used this to shed light on how replication accuracy can gradually increase through the acquisition of new catalytic functions. We reasoned that such a stepwise increase in the accuracy of template-guided ligation (or polymerization) could enable the faithful replication of longer and longer polymers over time and might, therefore, offer a solution to Eigen’s error threshold paradox (Eigen, 1971; Eigen and Schuster, 1978a, 1978b, 1977). This extended model, thus, explores the possibility that error-prone replicases could cooperate with mutation inhibitors to raise the viability of an entire polymer system and thereby elevate the maximum length of functional polymers that can be reliably replicated.

We used the basic and extended models to explore conditions conducive to the persistence of “functional” polymers. We considered variations in both external conditions (such as spatial structure and dilution rate) and internal factors (such as sequence length and catalytic efficiency). Our simulations suggest that spatially structured environments with connected compartments can result in multilevel selection that fosters the persistence of cooperating polymers. In line with expectations, we found that the viability of cooperative polymer systems is positively correlated to food flux and catalytic efficiency and is negatively correlated to the length of catalytic polymers, dilution rate, and mutation.

Together, our results suggest that, given diverse short polymers, an environment that promotes multilevel selection, and sufficient flux of resources that can react to produce activated monomers, new catalysts that arise can be maintained through the collective effects of multiple cooperating polymers. Moreover, our data suggest that cooperative systems can complexify in a stepwise and open-ended manner through the addition of new functions and/or through the discovery of longer sequences that confer the same functions but with higher catalytic efficiencies. Thus, our theoretical work provides useful clues for designing future *in silico* and *in vitro* experiments aimed at clarifying the pre-cellular emergence and complexification of genetic systems on the path to complex life.

## 2. Results

### 2.1. A simplified spatial model to efficiently simulate prebiotic nucleic acid chemistry

We set out to explore the ability of cooperative polymer systems to survive and evolve in a prebiotic environment using an abstract model that captures the effects of diverse physicochemical factors in a manner that resembles real chemistry. A core challenge in building such a model is the combinatorial explosion that most RNA-world models confront. Even if template-guided polymerization is treated as a single reaction (as in replicator equations (Schuster and Sigmund, 1983; Sigmund, 1986; Stadler, 1991)), and only two monomer species are considered, the computational load still increases exponentially with the maximum polymer length allowed. For example, for a maximum length of 20, there are more than 10^6^ single-stranded polymer species that need to be tracked. Moreover, if (partially) double-stranded polymers and the complexes formed by replicases, templates, and nascent strands are considered, the number of tracked species would be orders of magnitude higher still. Additionally, if we not only modelled tracked polarized addition (polymerization) or subtraction (exonuclease-like) of a single base at a time, which mimics RNA replication in living organisms, but also allowed random breakage of polymers and ligation among fragments of any length, which could be realistic in prebiotic environments, then the number tracked reactions would quickly become infeasible.

Inspired by Polymerase Chain Reaction (PCR) and the classical computer game Tetris®, we developed a model that significantly simplifies the process of simulating polymer systems and avoids the need to pre-specify maximum sequence length. We then explored an initial case of cooperation between a “kinase,” which catalyzes the formation of activated monomers from food, and a “ligase,” which links activated monomers or polymers to other monomers or already formed polymers. Here, the two catalysts cooperate to convert food into more copies of themselves and their reverse complements as well as other non-functional sequences.

Nevertheless, providing service to the entire polymer system incurs a cost: when a sequence is acting as a catalyst, it cannot act as a template at the same time. An extended model added a third function that results in the inhibition of mutation, but we will postpone describing this model until a later section.

In our model, molecules can be adsorbed to the bottom surface of a reactor; an adsorbed molecule may or may not have polarity along the horizontal dimension. Polymers are built from activated monomers A and B that have polarity and the capacity for reverse-complementary base-pairing, analogous to nucleotide triphosphates. Kinase polymers catalyze the reversible reactions E + F ⇌ A + W and E + G ⇌ B + W, which synthesize activated free monomers A and B from energetic food (E) and structural foods (F, G). Activated monomers (A, B) and structural food (F, G) are polar, meaning that the two borders of sites that they occupy have different possible reactions. Energetic food and waste, in contrast, are nonpolar. Kinase activity generates the building blocks of catalytic and genetic polymers, as in metabolism.

To enable mutation, we allow for A and B to acquire a “wildcard” status, M or N, respectively, which allows them to pair equally well with any monomer types. Such a mode of mutation resembles chemical changes to nucleic acids, such as certain methylation or amination reactions, which have the potential to be correctly or incorrectly repaired.

Ligase activity provides the basis for genetic encoding insofar as ligase catalysis is sensitive to base-pairing with a template sequence, meaning that there is a tendency for genetic information to pass from a template to a complementary strand (Tkachenko and Maslov, 2018). In our model, a ligase can link monomers regardless of whether they are free or incorporated in a polymer; for example, a ligase would act on reactions such as ABAA + B ⇌ ABAAB and ABA + AB ⇌ ABAAB.

To simulate templated-guided synthesis of polymers, our model considers a reactor of which the bottom surface is narrow enough such that it can be viewed as a quasi-1D line. This linear bottom surface can adsorb two layers of molecules (like the game Tetris®), leaving other molecules (if any) moving around in the solution above these layers. The model cycles through three phases: adsorption, reaction, and resuspension (Fig. 1). Because kinase-catalyzed and ligase-catalyzed reactions only occur at the borders between adsorbed molecules, the possibility of a reaction occurring depends only on the occupants of adjacent surface sites (Fig. 1(b)). As a result, during the reaction phase, the model simply needs to track and update the states of the borders where reactions may occur, meaning that it is not necessary to track individual polymer sequences. More details of the model can be found in Supplemental Materials & Methods.

**Fig. 1.**
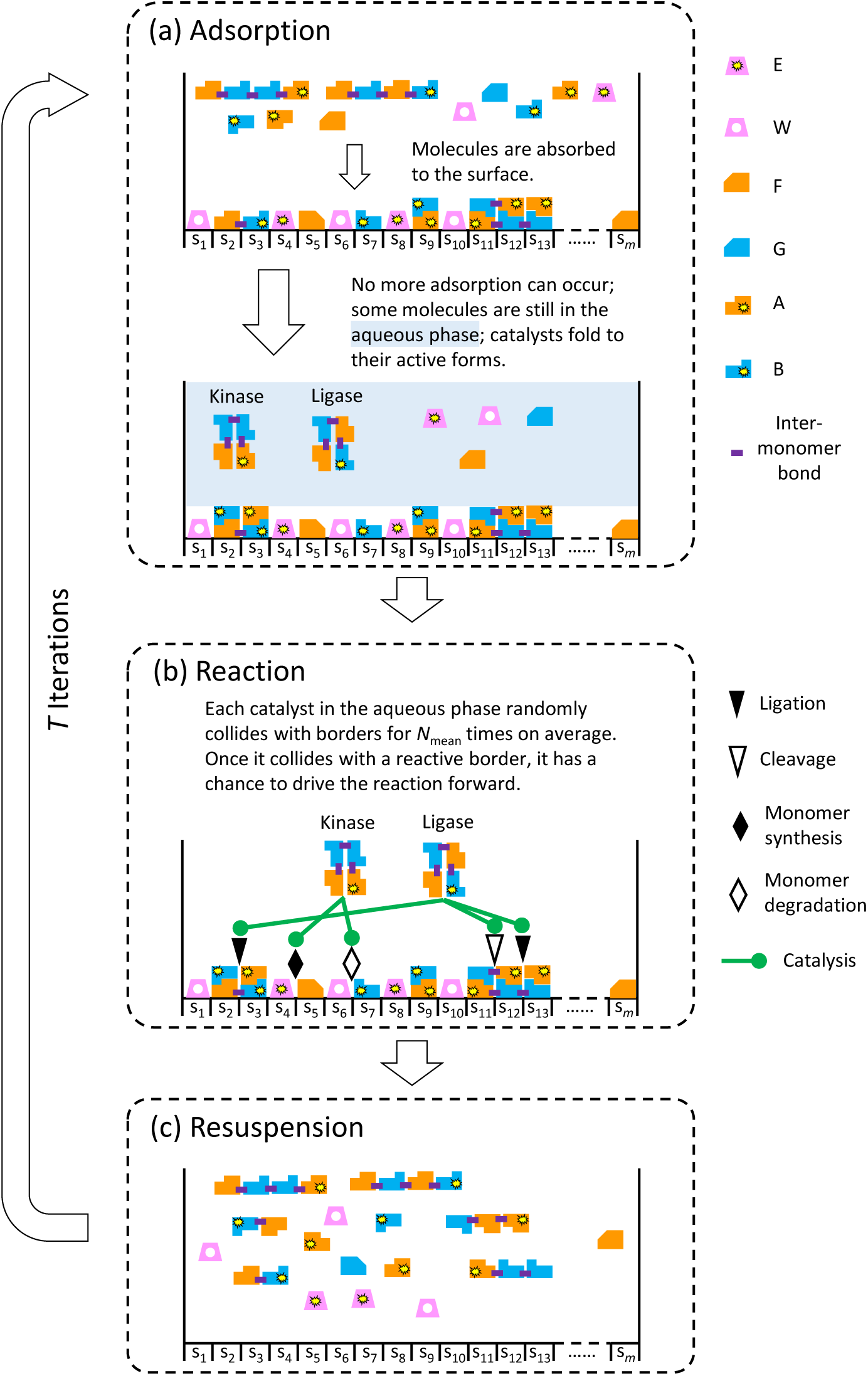
Three phases of the basic model. Each reactor has *m* sites. Each site can adsorb a food molecule, a waste molecule, a free monomer, or an incorporated monomer. Energy food (E) and waste (W) are nonpolar, whereas structural foods (F and G), free monomers (A and B), and polymers are polar. **(a)** Phase 1: Adsorption. First, free and incorporated monomers stochastically become “wildcards,” which are monomers that can pair with any type of monomer. The fraction of monomers that are converted into wildcards correlates with the probability of a mismatch mutation occurring during ligation. Second, molecules are randomly adsorbed to the surface until no more adsorption can occur. When a polar unit, such as a free or incorporated monomer, is adsorbed to the surface, it stochastically makes its adjacent sites inaccessible to other polar molecules, which lowers the chances of fusion mutations. **(b)** Phase 2: Reaction. Each catalyst in the aqueous phase randomly collides with borders between adjacent surface sites *N* times, where *N* is drawn from a Poisson distribution. Once a catalyst collides with a reactive border, it has a chance to cause a reaction. In this phase, wildcard monomers behave the same as normal monomers. **(c)** Phase 3: Resuspension. All adsorbed molecules are released to the aqueous phase and any remaining wildcard monomers are converted back to normal monomers.

In Phase 1 (adsorption), free and incorporated monomers have a probability of becoming wildcards. Then, polymers, free monomers, energetic and structural foods, and waste are adsorbed to the surface until no more adsorption can occur (Fig. 1(a)). Any free monomers or polymers that are not adsorbed in the first layer can be adsorbed onto a second layer, similar to the way that a Tetris® brick can fall on another brick (Fig. 1(a)). The second-layer adsorption follows reverse-complementary base-pairing rules and continues until no more adsorption can occur. After this phase, molecules or incorporated monomers that occupy adjacent sites determine whether the borders between these sites are “reactive borders” (Fig. 1(b)), where a bond may be formed or broken during the reaction phase.

In Phase 2 (reaction), catalytic polymers that were not adsorbed are allowed to randomly collide with borders (Fig. 1(b)). Whenever a catalytic polymer (e.g., ligase) collides with a reactive border, it has a probability, its *catalytic efficiency*, of causing a reaction to occur. For example, a reaction might entail two unconnected monomers becoming connected (Fig. 1(b), black triangle), a polymer being split (Fig. 1(b), empty triangle), or a structural food reacting with an energetic food to generate a free monomer and waste (Fig. 1(b), black diamond). For a catalytic polymer, the number of collisions with borders within a reaction phase follows a Poisson distribution and negatively correlated to the polymer mass (reflecting the slower diffusion of larger molecules). Spontaneous reactions can be implemented by assuming that natural catalysts are present at some baseline concentration. For more details, see Supplemental Materials & Methods.

The number of reaction events that occur in the reaction phase is mainly determined by the number of catalytic polymers and the number of reactive borders. Under this approach there is no need to re-calculate reaction propensities within an iteration since the number of catalysts and reactive borders does not change and there are only two general types of reactive border: (i) the borders between nonpolar molecules (E, W) and properly oriented polar foods (F, G) or properly oriented free monomers (A, B) in the first layer, which may undergo the conversion of energetic and structural foods into free monomers and waste or the reverse reaction, and (ii) the borders between similarly-oriented monomers in the second layer, which may undergo the formation or breakage of bonds between monomers. The strategy of only tracking reactive borders circumvents the combinatorial explosion problem, and can be thought of as a case where catalysts are shortsighted, and only care about whether a border on the surface can be attacked, regardless of what molecules are consumed or produced after a bond is formed or broken. As a result, reactions that are catalyzed by the same type of catalysts are not independently tracked but lumped together and only the sites next to reactive borders have their statuses tracked, which dramatically decreases the computational load.

In Phase 3 (resuspension), all adsorbed molecules are released (Fig. 1(c)), returning the system to the original configuration with a different set of molecules. Given that at least some kinase and ligase polymers exist at the beginning, iterating these three phases – adsorption, reaction, and resuspension – for multiple cycles is conceptually similar to iterating the steps of annealing, extension, and denaturation in PCR. Multiple such cycles form a generation (Fig. 1).

To allow for multilevel selection, which could aid in the enrichment of cooperating polymers (Hogeweg and Takeuchi, 2003; Szathmáry and Demeter, 1987; Takeuchi and Hogeweg, 2009), we created a circular array of reactors (Fig. 2(a), where the leftmost and rightmost reactors are also neighbors). After a generation ends, a fraction of each reactor’s contents are replaced by food from an external source (Fig. 2(b)), and then reactors exchange a portion of their contents, representing overflow or diffusion (Fig. 2(c)). The proportion of the contents that are replaced each generation is the *dilution rate*.

**Fig. 2.**
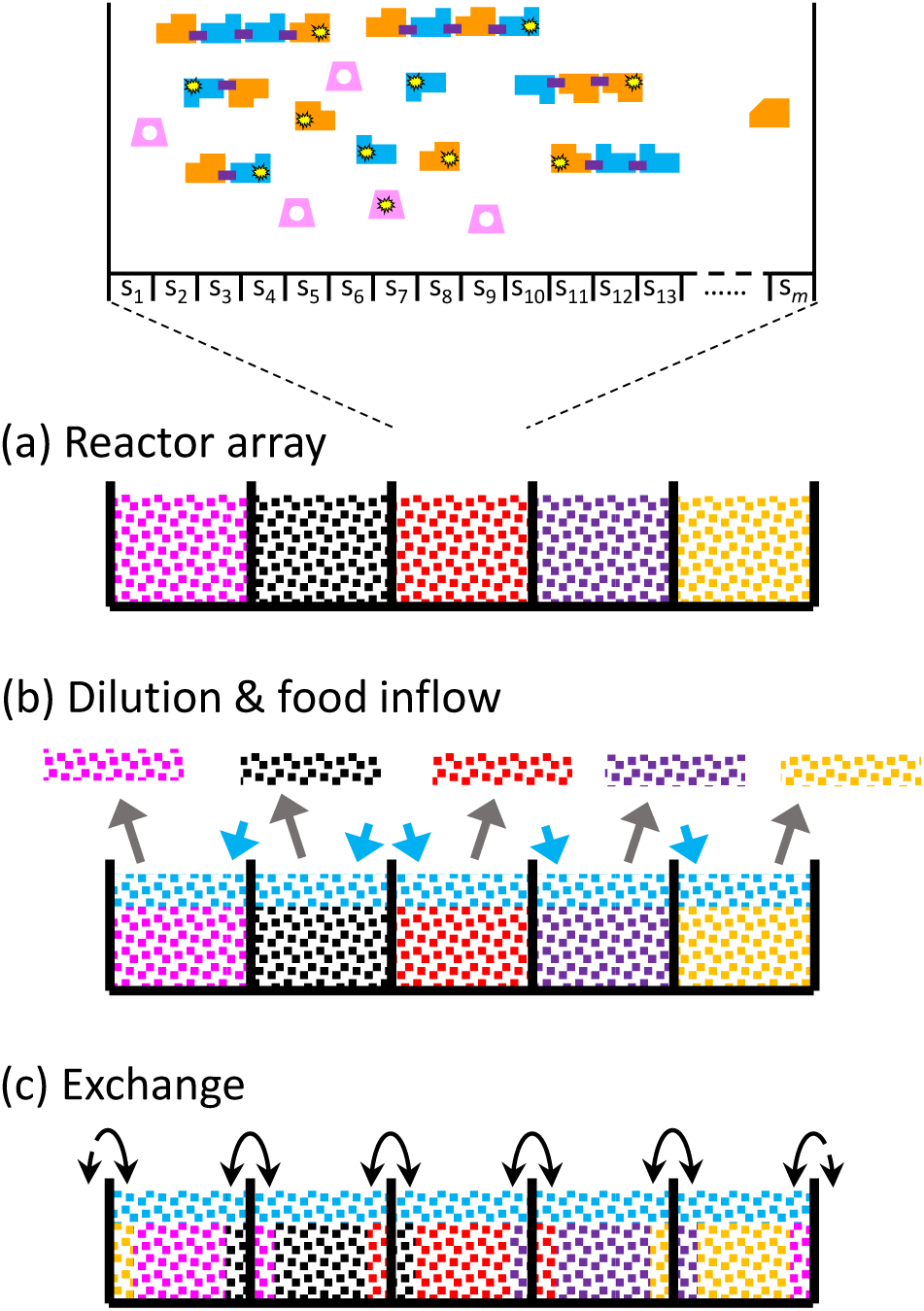
A reactor array. **(a)** Multiple reactors can form a circular array, where the rightmost and the leftmost reactors are neighbors. **(b)** Reactors have a portion of the contents (grey arrows) replaced by foods (cyan arrows). **(c)** Reactors exchange their contents. This example shows the neighborhood-dispersal regime; there may also be other schemes; see Section 2.2 and Fig. 3.

To demonstrate the ability of this model to handle long polymers, we initiated the reaction system with only food species (E, F, and G), running the simulations with high levels of spontaneous activation/ligation and an elevated probability of spontaneous fusion (see Supplemental Materials & Methods). We ran 30 replicates, each with a single reactor of 200 sites (i.e., 199 borders), for 20 generations. We found that the longest polymers present in the final populations have an average length of ∼44 (Table S1) and the longest polymer among all replicate simulations is a 63-mer (Table S1, Replicate 3). Running all the 30 replicates in parallel took approximately 29 hours using the high throughput computing service at the Center of High Throughput Computing at the University of Wisconsin–Madison (Center for High Throughput Computing, 2006).

### 2.2. Externally provided compartments and slow inter-compartment exchange favor cooperating polymers’ persistence

Inspired by classical research on the evolution of cooperation driven by group selection (Smith, 1964), we hypothesized that externally provided spatial structure, such as pores in porous rocks, combined with slow inter-compartment exchange, could promote cooperation and suppress cheating. Here, cheating refers to the phenomenon that polymers that are neither catalysts providing system-level service nor reverse complements of these catalysts compete with the catalysts and their reverse complements for the chances of being templates. To test our hypothesis, we evaluated a scenario in which synthesis of polymers requires the presence of a kinase polymer that promotes the activation of nucleotides and a ligase polymer that converts monomers into polymers in a template-guided manner, and asked how likely it is that both the kinase and ligase functions would persist if they happened to both arise spontaneously in a single reactor. To test our hypothesis, we performed comparisons across three groups of simulations.

The first group experienced *neighborhood-dispersal*, where 30 reactors (each with 100 sites) are connected to form an array, and a portion of each reactor’s volume is exchanged with its two neighbors (Fig. 3(a)). The second group of 30 reactors (each with 100 sites) experienced *global-dispersal*, which resembles Maynard Smith’s haystack model (Smith, 1964), where all reactors release a portion of their volume to a shared pool, and then molecules from this pool are randomly redistributed among all reactors (Fig. 3(b)). The third *well-mixed* group entailed a single large reactor, with 3000 sites (Fig. 3(c)).

**Fig. 3.**
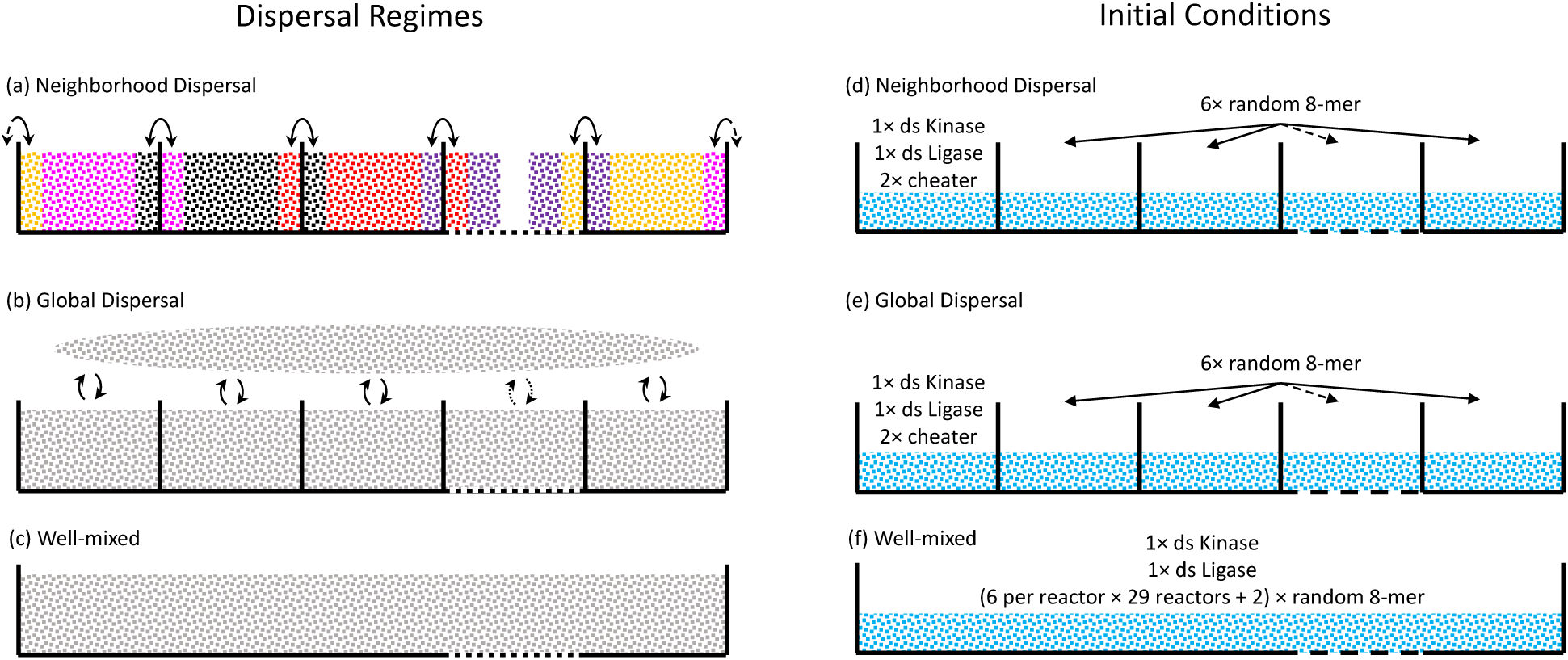
Three dispersal regimes and their initial conditions. **(a)** In the neighborhood-dispersal regime, a randomly selected 60% of a reactor’s contents (by molecular count) remain in the same reactor, 20% move to the left reactor, and 20% move to the right reactor. **(b)** In the global-dispersal regime, a randomly selected 60% of a reactor’s contents remain in the same reactor and 40% contribute to a shared pool that is then randomly redistributed among all the reactors. **(c)** In the well-mixed regime, there is only one giant well-mixed reactor. **(d)(e)** Simulations for the neighborhood-dispersal and global-dispersal groups are initialized with one 8-mer kinase (ABAABABB) and its reverse complement (AABABBAB), one 8-mer ligase (BBAAABAB) and its reverse complement (ABABBBAA), and two random 8-mer cheater polymers in the first reactor, whereas the other 29 reactors are seeded with six random 8-mers. **(f)** Simulations for the well-mixed group are initialized with the same sequences as the other two conditions, but all in a single reactor.

All simulations used low spontaneous reaction rates and low probabilities of fusion-induced mutations and mismatch-induced mutations (see Supplemental Materials & Methods). The neighborhood-dispersal and global-dispersal groups were initialized with one reactor seeded with one 8-mer kinase (ABAABABB) and its reverse complement (AABABBAB), one 8-mer ligase (BBAAABAB) and its reverse complement (ABABBBAA), and two 8-mer cheaters, while the other 29 reactors were each seeded with six random 8-mers. All reactors began with their food concentrations the same as the external food source (Fig. 3(d-e)). The well-mixed group included the same 180 sequences (6 polymers times 30 reactors) all in the single reactor (Fig. 3(f)).

For each group, multiple simulations were performed, which allowed us to estimate the proportion of simulations where both cooperators persisted. We defined persistence as there being, after 50 generations, at least one reactor where the kinase, the ligase, and their reverse complements were all present.

Although we showed in Section 2.1 that our model can handle polymers longer than 60-mers, we chose to focus initially on 8-mer catalysts. We do not need long functional polymers to assess the relative effects of different factors on their persistence since, even with 8-mers, there are many more non-functional cheater sequences ((2^2^ + 2^3^ + … + 2^8^) – 4= 504) than functional sequences (2) and reverse complements of functional sequences (2). By using sequences of this length, computation is more efficient, allowing us to explore many different factors within a reasonable timeframe.

As shown in Fig. 4, the probability of persistence is sensitive to catalytic efficiency, and is enhanced in the neighborhood-and global-dispersal groups, suggesting a role for multilevel selection. When catalytic efficiencies are low (kinase efficiency = 0.0005, ligase efficiency = 0.02), the cooperating catalysts are only ever viable in the neighborhood-dispersal group (Fig. 4(a)). In most cases where persistence did not occur, extinction occurred in the first few generations. The latter helps explain why the neighborhood model is especially conducive to persistence: copies of the functional species produced in the first generation only spread out to its two neighbors, increasing the chance that one or both neighbors will be initiated in the second generation with both cooperating catalysts, while the cheater sequences in other reactors have to wait for longer time before they have access to functional polymers or die out. When catalytic efficiencies are high (kinase efficiency = 0.005, ligase efficiency = 0.2), catalysts could persist in all three cases (Fig. 4(b)), but even here the persistence probability for well-mixed group was significantly lower than the other two.

**Fig. 4.**
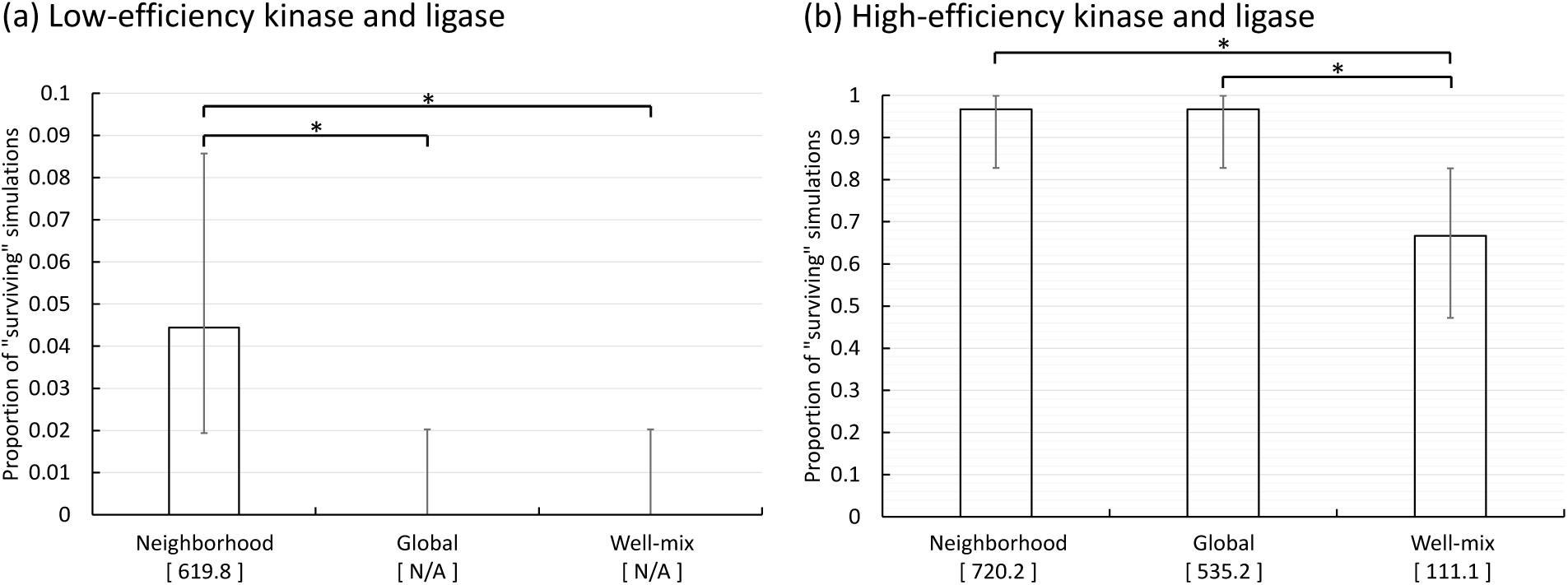
Compartmentalization and slow inter-reactor exchange help cooperating polymers survive. Graphs show the proportion of simulations where the kinase and ligase sequences both survived after 50 generations. Error bars show the 95% Clopper-Pearson confidence intervals. The number in brackets below a column indicates the summed count of catalytic sequences and their reverse complements per surviving simulation at the end. **(a)** Each column is based on 180 replicate simulations; kinase efficiency = 0.0005, ligase efficiency = 0.02. **(b)** Each column is based on 30 replicate simulations; kinase efficiency = 0.005, ligase efficiency = 0.2. Note that **(a)** and **(b)** have different vertical axis scales. For all simulations, the dilution rate was 0.05; external food counts per unit volume were 1000 for E, 500 for F, and 500 for G. Significant differences are indicated by stars.

Since the neighborhood-dispersal model is geologically plausible and seems to be the most effective organization for enabling the persistence of cooperating polymers in this case, we utilized this model for all subsequent analyses.

### 2.3. High food concentration and low dilution rates help cooperating polymers survive

It has been shown using *in silico* continuously stirred flow tank reactors, that the carrying capacity of the members of an autocatalytic cycle is positively correlated with the concentration of food in the influx and negatively correlated with the dilution rate (Peng et al., 2020). Since molecules subject to template-mediated replication, such as the nucleic acid molecules modelled here, are members of autocatalytic cycles, we wondered whether similar results would hold for such a more complex, multi-autocatalytic-cycle system.

Defining the combination of high-efficiency kinase and ligase as our starting point, we assessed how system viability changes when food concentration decreases and dilution rate increases. As expected, lower food concentrations and higher dilution rate are associated with lower viability (Fig. 5).

**Fig. 5.**
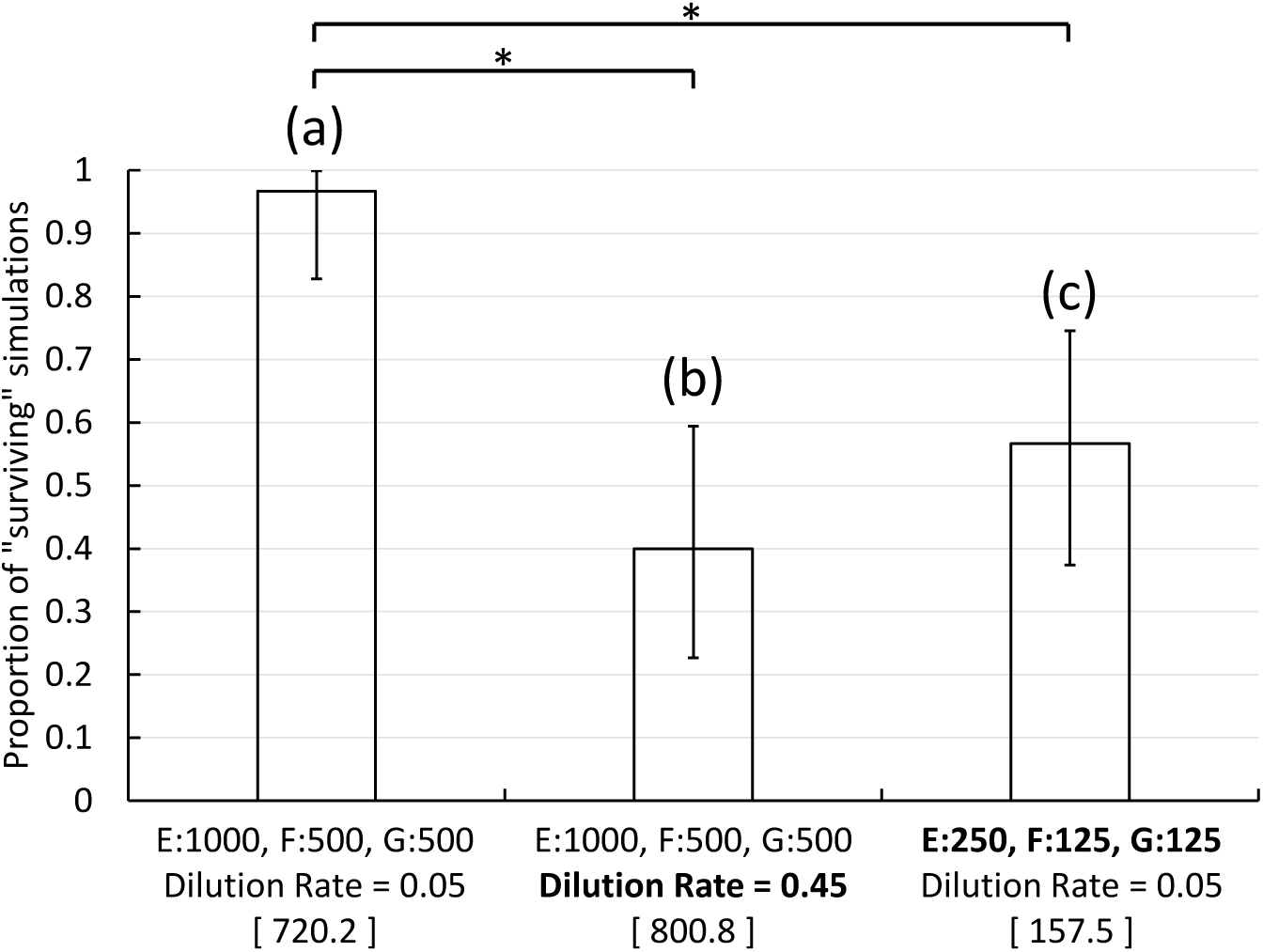
A higher food concentration and a lower dilution rate are associated with elevated viability. The **(a)** column is directly copied from the column labeled “Neighborhood” in Fig. 4(b) and serves as the reference point. **(b)** Dilution rate was increased to 0.45. **(c)** External food counts per unit volume were decreased to E:250, F:125, G:125. Error bars show the 95% Clopper-Pearson confidence intervals. The bracketed number below a column indicates the summed count of catalytic sequences and their reverse complements per surviving simulation at the end. Each column is based on 30 replicate simulations. Significant differences are indicated by stars.

### 2.4. Systems consisting of polymers with higher catalytic efficiencies are more likely to survive

The data summarized in Fig. 4 already showed that elevating the catalytic efficiency of both the kinase and ligase polymers increases the probability of persistence. We wondered whether a similar benefit might be seen if just one of the functional catalysts is more efficient, since this might enable ratchetting up of system viability, one function at a time. Indeed, simulations using different combinations of high- or low-efficiency kinase and high- or low-efficiency ligase (Fig. 6) show that enhancing either catalytic efficiency ten-fold significantly enhances the mutual persistence probability.

**Fig. 6.**
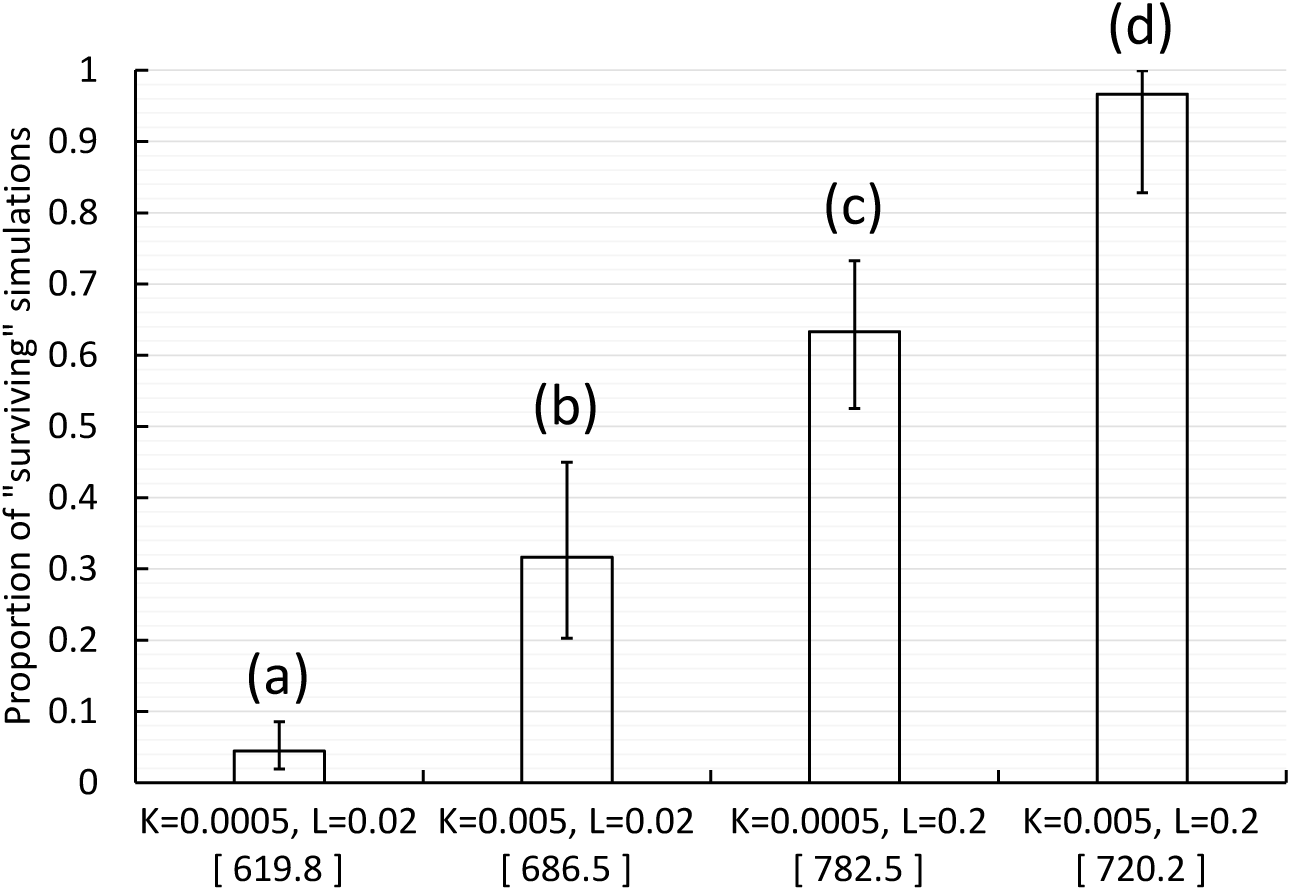
Catalytic efficiencies of both functions are positively correlated to viability. Error bars show the 95% Clopper-Pearson confidence intervals (all columns are significantly different). The bracketed number below a column indicates the summed count of catalytic sequences and their reverse complements per surviving simulation at the end. **(a)** Kinase efficiency = 0.0005, ligase efficiency = 0.02, 180 replicates. **(b)** Kinase efficiency = 0.005, ligase efficiency = 0.02, 60 replicates. **(c)** Kinase efficiency = 0.0005, ligase efficiency = 0.2, 90 replicates. **(d)** Kinase efficiency = 0.005, ligase efficiency = 0.2, 30 replicates. Except as indicated, initial conditions and model parameters are the same as Fig. 4.

These results are easy to explain, since higher catalytic efficiencies bring about more conversion of food into polymers within a generation (Fig. 1), which means that catalytic polymers (of both kinds) are less likely to be lost during inter-generation dilution (Fig. 2). In this case, parameter values appear to fall within a range such that survival is sensitive to both kinase and ligase efficiency; however, with these parameter values, the system seems to gain more from a tenfold increase in ligase activity than kinase.

### 2.5. Systems consisting of shorter polymers are more likely to survive

Longer polymers have greater potential to adopt diverse secondary and tertiary structures, opening up more potential catalytic roles and probably raising the maximum plausible catalytic efficiency. For example, long polymers have more potential for conformational change, which can greatly increase catalytic efficiency (Rivoire, 2024). However, the rate at which longer functional polymers can be produced is lowered by the fact that their production consumes more resources and requires more reactions. Additionally, if mutations are frequent, longer templates will tend to yield more non-functional products.

To assess the influence of polymer length on the viability, we used the combination of low-efficiency kinase and high-efficiency ligase as the reference, as shown in Fig. 6 column (c). We ran three additional groups of simulations where catalytic efficiencies were unchanged, but the length of the ligase and/or kinase was increased to 10 monomers by adding an A monomer and a B monomer flanking the original 8-mers. The results (Fig. 7) show that the combination of 8-mer kinase and 10-mer ligase (Fig. 7 column (b)) or 10-mer kinase and 8-mer ligase (Fig. 7 column (c)) have similar viabilities, and both are lower than the viability of the combination of 8-mer catalysts (Fig. 7 column (a)) and higher than the combination of two 10-mer catalysts (Fig. 7 column (d)). These results show that holding other properties similar, a system relying on longer catalytic polymers has lower viability.

**Fig. 7.**
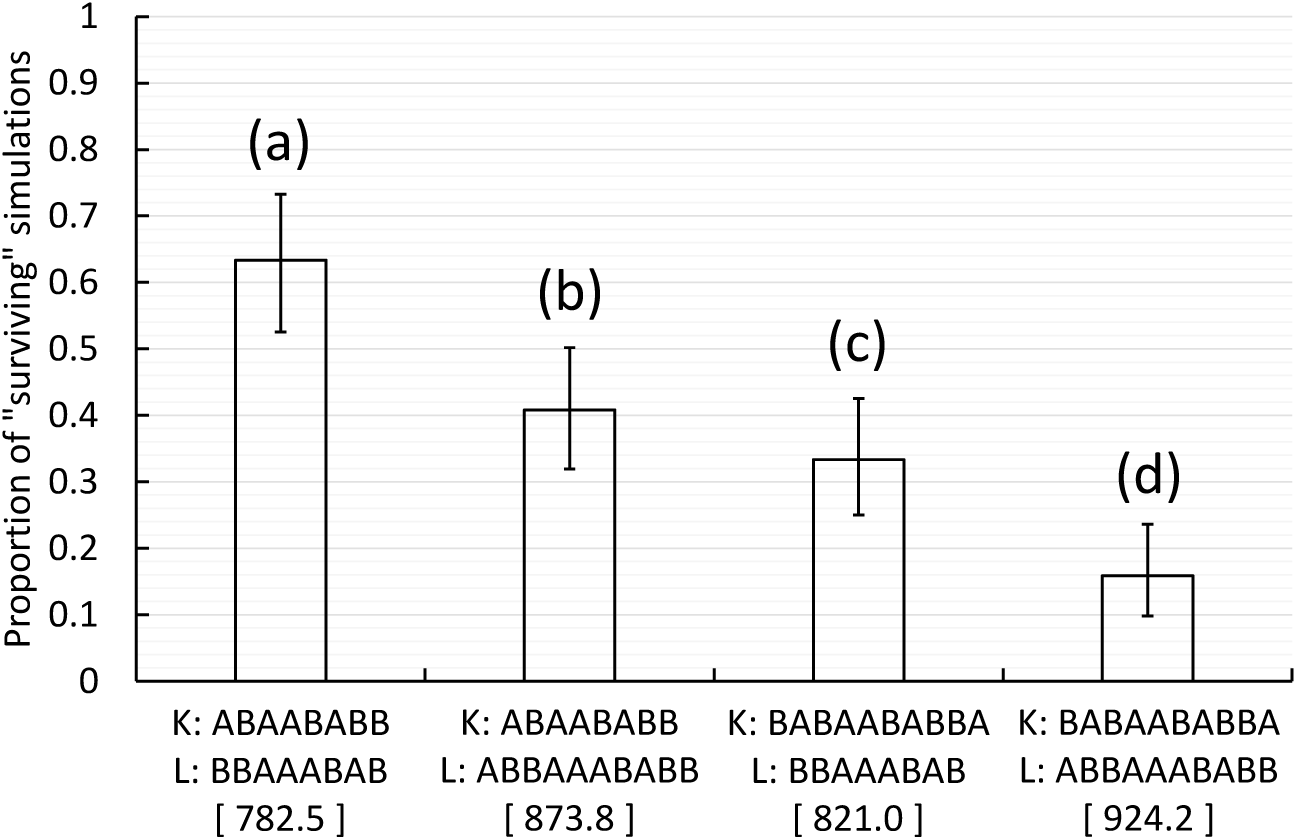
Catalyst length is negatively correlated to viabilities of polymer systems. Initial conditions and model parameters are the same as Fig. 6(c) except for the sequences of catalytic polymers (kinase efficiency = 0.0005, ligase efficiency = 0.2). Error bars show the 95% Clopper-Pearson confidence intervals. The bracketed number below a column indicates the summed count of catalytic sequences and their reverse complements per surviving simulation at the end. **(a)** Kinase: ABAABABB, ligase: BBAAABAB, 90 replicates. **(b)** Kinase: ABAABABB, ligase: ABBAAABABB, 120 replicates. **(c)** Kinase: BABAABABBA, ligase: BBAAABAB, 120 replicates. **(d)** Kinase: BABAABABBA, ligase: ABBAAABABB, 120 replicates. Apart from columns (b) and (c), all other pairs of columns have significantly different survival probabilities.

It is worth noting that by comparing the probability of survival of low-efficiency 8-mer kinase with high-efficiency 10-mer ligase (∼41%; Fig. 7 column (b)) to that of low-efficiency 8-mer kinase with low-efficiency 8-mer ligase (∼4%; Fig. 6 column (a)), we see that a higher catalytic efficiency can compensate for the cost of longer polymers.

### 2.6. Systems with higher fidelity of replication are more likely to survive

It has long been appreciated that early template-guided replication mechanisms in the prebiotic world likely had high error rates, which would make it difficult to reliably replicate sequences beyond some maximum length: Eigen’s error threshold (Eigen, 1971; Eigen and Schuster, 1978a, 1978b, 1977). It is generally believed that some kind of proofreading would be needed to access the polymer space beyond this length threshold. However, paradoxically, adding a proofreading function to a polymerase or ligase RNA likely requires adding additional functional motifs, and thus additional length, to the molecule. We wanted to explore whether this paradox could be resolved by the appearance of an independent error-inhibiting polymer that is not polymerase or ligase but rather integrated into an ecosystem of cooperating functional polymers in such a way as to allow longer polymers to be reliably replicated. Such an ecosystem could, we reasoned, allow for progressively longer catalysts, perhaps allowing for the eventual emergence of proofreading replicases.

To explore this question, we needed to introduce a mechanism of mutation that could be counteracted by a plausible, and easily modelled, error inhibition mechanism. As shown in Fig. 1 and described in more detail in Supplemental Materials and Methods, free and incorporated monomers were assigned a probability of becoming wildcards at the beginning of the adsorption phase. Wildcards are modifications to the A or B monomers that make those monomers equally likely to match any monomers during adsorption. By including wildcards, we allow for mismatch-induced mutations.

To implement an error inhibition mechanism, we inserted a stage immediately following the introduction of wildcards: wildcard correction. This is where wildcards have the chance to be converted back into normal monomers due to the action of a catalytic *mutation inhibitor*. The model we implemented assumed that during this wildcard correction stage, the more mutation inhibitor polymers in the solution, the higher the probability that a given wildcard is converted back to its original monomer type, and if there is no mutation inhibitor present, wildcards are never converted back to normal monomers before the reaction phase. The efficiency of a mutation inhibitor is described by its correction efficiency index *v* ∈ (0, 1) (see Supplemental Materials and Methods), such that for inhibitor species with different *v*’s but the same count, the one with a higher *v* has a larger probability of converting a given wildcard back to a normal monomer. Note that the frequency of wildcards after the correction stage does not directly correspond to the mutation rate (in mutations per monomer per cycle), because a wildcard does not necessarily lead to mismatch (e.g., a wildcard of A happens to match a normal B) and a mismatch does not necessarily lead to a mutation (e.g., the first-layer template happens to be a single free monomer). Nonetheless, the realized mutation rate should relate positively (and monotonically) to the wildcard frequency and, thus, negatively to the efficiency of wildcard correction.

With such an extended model, it becomes possible to assess how mutations interact with polymer length (as a rough proxy for molecular complexity) and catalytic efficiency to influence the viability of polymer systems. We considered cases of pairs of high-efficiency 8- or 10-mer kinases and ligases. The analyses were similar to the cases seen in Fig. 6 column (d) and Fig. 7 column (d), except that we raised the efficiency of the 10-mer kinase to 0.005 and simulations were initiated with the first reactor containing not only the ligase and kinase but also a specific 8-mer sequence (ABBABBAA), which will be defined as the mutation inhibitor in later analyses, and its reverse complement (ABBABBAA). We then explored wildcard-introduction probabilities from 0.001 to 0.1 (as contrasted with a rate of 10^-6^ in prior simulations).

Fig. 8 confirms that the viability of a polymer system is negatively correlated with the wildcard-introduction probability and, thus, mutation rate. Moreover, a system relying on catalytic 10-mers is more sensitive to mutations than a system relying on catalytic 8-mers. It is noteworthy, however, that systems can persist even with a wildcard-introduction probability as high as 2%, presumably because, if catalytic efficiencies are high enough, polymer systems can synthesize enough correct copies to compensate for the many non-functional descendants that are generated.

**Fig. 8.**
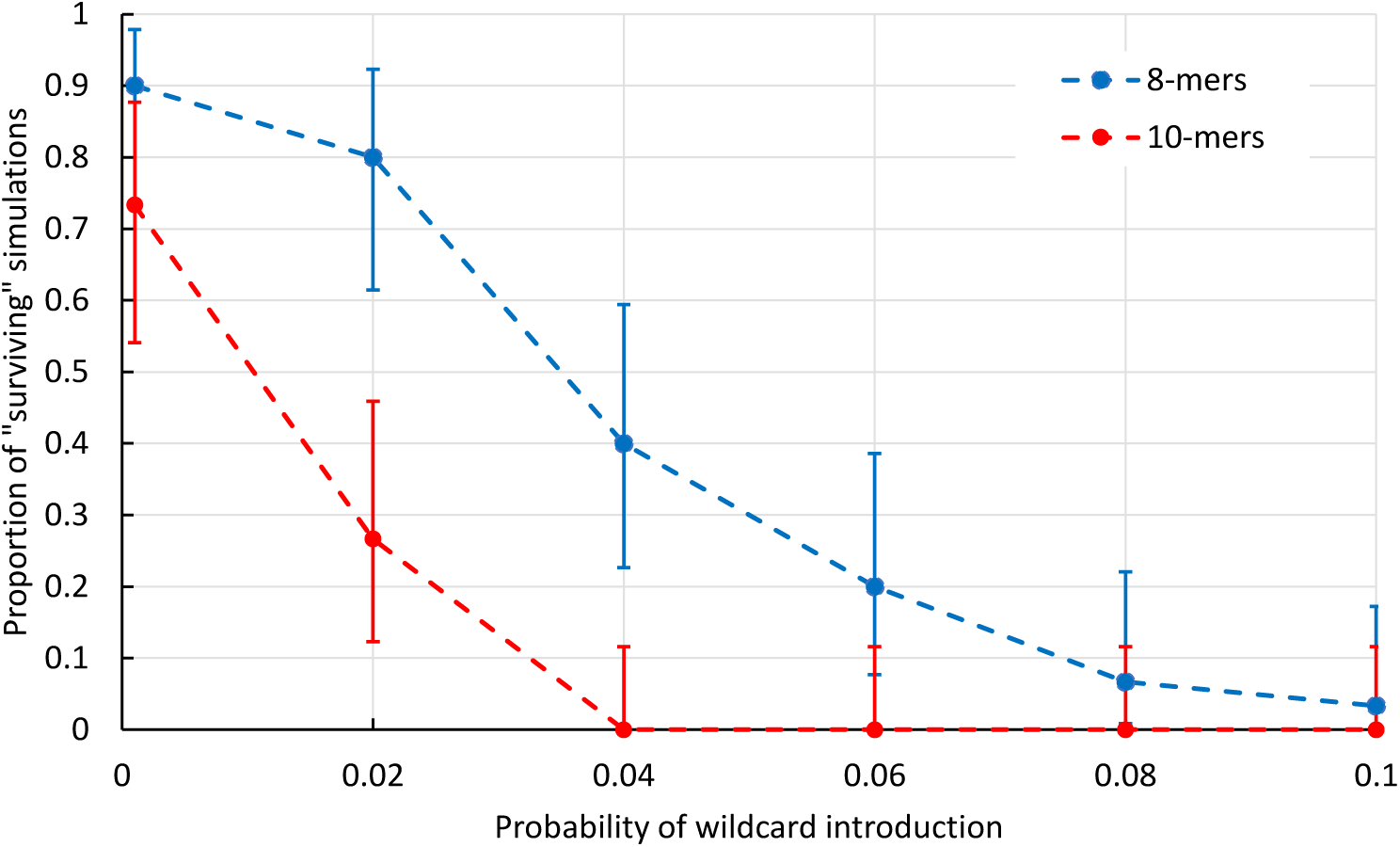
Interaction between mutation propensity and catalyst length influences viabilities of polymer systems. Error bars show the 95% Clopper-Pearson confidence intervals. Probability of wildcard introduction per adsorption phase varies between 0.001 and 0.1. All data points have 30 replicate simulations. **Blue series:** Kinase: ABAABABB, ligase: BBAAABAB. **Red series:** Kinase: BABAABABBA, ligase: ABBAAABABB. Except as stated, model parameters were the same as Fig. 6(d).

To see if a mutation inhibitor catalyst can rescue the low viability of the 10-mer cooperative system with a wildcard-introduction probability of 0.04 (Fig. 8), we ran another series of simulations over a range of catalytic efficiencies. The index measuring the efficiency of the mutation inhibitor varied from 0 to 0.9. Note that since the new kinase-ligase-inhibitor cooperative system had three instead of two functions, the viability was measured by the proportion of simulations where all three functional polymers (and their reverse complements) were present in at least one reactor at the end of simulation. As expected, the mutation inhibitor greatly increases the viability of the polymer system (Fig. 9). Interestingly, the survival probability seen in this system, whose kinase and ligase are 10-mers, can equal or even exceed the level seen in the 8-mer cooperative system with a wildcard-introduction probability of 0.04 (compare to the blue line in Fig. 8).

**Fig. 9.**
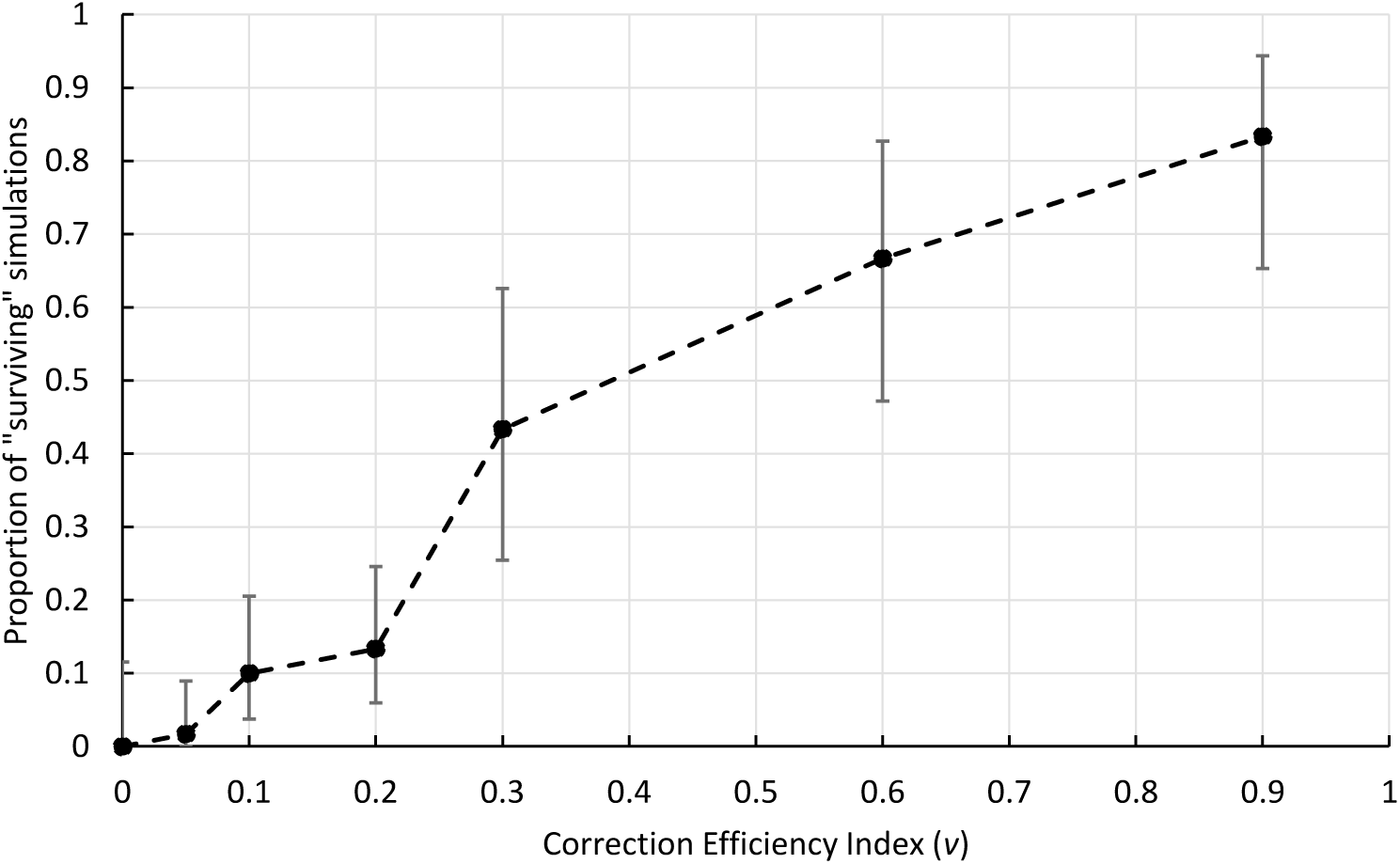
Mutation inhibitor rescues the viability of a cooperative polymer system. Model parameters are the same as the red data point with wildcard-introduction probability being 0.04 in Fig. 8, except that the efficiency of the inhibitor ABBABBAA was varied from 0 to 0.9. Error bars show the 95% Clopper-Pearson confidence intervals. For *v* ∈ {0, 0.3, 0.6, 0.9} there are 30 replicate simulations; for *v* ∈ {0.05, 0.1, 0.2} there are 60 replicate simulations. Viability (vertical axis) is measured by the proportion of simulations where kinase BABAABABBA, ligase ABBAAABABB, mutation inhibitor ABBABBAA, and their reverse complements are all present in at least one reactor at the end of the simulation.

## 3. Discussion

In this study, we developed an abstract model of prebiotic chemical ecosystems with template-guided replication and used it to explore cooperative persistence among catalytic polymers. The model allowed us to show that the viability of cooperative polymer systems is enhanced by the existence of a spatial structure conducive to multilevel selection. This result aligns with prior studies showing that such multilevel selection can promote cooperative behaviors in modern organisms (Buttery et al., 2012; Van Dyken et al., 2013). The importance of multilevel selection in origins of life, and especially to control cheaters, has previously been demonstrated using both computer simulations (Shay et al., 2015; Takeuchi and Hogeweg, 2009) and empirical experiments (Matsumura et al., 2016; Mizuuchi and Ichihashi, 2018). Although theoretical work has suggested that intrinsic compartmentalization, such as conferred by protocells, is even more effective than the spatial structure and extrinsic compartmentalization studied here (Shah et al., 2019), it is challenging to explain the emergence of growing and dividing protocells containing cooperating polymers in the absence of prior multilevel selection. In contrast, it is easy to imagine situations that would approximate an array or environmentally delimited ecosystems, as studied here, which suggests that polymer cooperation could long precede cellular compartmentalization. In addition, we showed that intrinsic physicochemical properties of polymers, such as catalytic efficiencies and length, affect the viability of cooperative polymer systems, which allows natural selection to potentially drive adaptive evolution of such prebiotic multi-molecular systems (Tkachenko and Maslov, 2018).

Our results, taken together, suggest a possible mechanism by which the genetic system of modern life could have emerged and overcome Eigen’s error threshold paradox (Eigen, 1971; Eigen and Schuster, 1978a, 1978b, 1977). Specifically, it seems plausible that ecosystems of short ribozymes catalyzing diverse, complementary functions, such as the synthesis of activated nucleotides (metabolic catalysts, kinases, etc.), RNA replication (e.g., polymerases, ligases), and mutation inhibition, could have emerged in conditions conducive to multilevel selection. In such settings, gradual improvements in the production of monomers, increases in the efficiency of their polymerization, and gradual increases in replication fidelity could gradually accumulate and, as a result, progressively longer (and potentially more efficient) ribozymes could become viable. This stepwise evolution could entail new catalysts that fulfill novel functions as well as ones that fulfill pre-existing functions, but with greater efficiency. Among other catalytic functions, it is easy to imagine that such progressive complexification could result in the eventual emergence of proofreading polymerases.

The strategy we pursued here was to presume the existence of simple catalytic polymers and investigate conditions conducive to their survival. While useful, this approach does not address the initial source of genetic diversity. One possibility is that there was sufficient non-enzymatic spontaneous synthesis and ligation for the shortest functional polymers, and their reverse-complements, to emerge by chance. However, when spontaneous synthesis is already efficient, functional polymers would have to have high catalytic efficiencies to significantly improve their survival probability. Alternatively, we can imagine two classes of environments. The first, represented by data shown in Table S1, would be one that promotes the spontaneous synthesis of polymers but lacks strong selection for catalytic activity. The second environment would be one that is unsuited to high rates of non-enzymatic polymer formation, perhaps because of the rapidity of wet-dry cycling, or a lack of catalytic minerals, but receives occasional input from the first type of environment. In such a situation, polymer dispersal from the former to the latter class of environments would set up the kinds of situations modelled here, where catalytic polymers appear periodically, but can only persist when they cooperate effectively with other polymers in those ecosystems.

Our results, especially Section 2.6, demonstrate that this model is flexible, and can be extended to explore diverse intrinsic and extrinsic factors. A few possible extensions are worth mentioning. We used a 1D circular array of reactors, but many real-world settings, such as mineral surfaces, porous rocks, hydrothermal fields/vents, or periodically wetting-and-drying shorelines, would be better represented by 2D (or even 3D) reactor arrays. It would be informative, therefore, to examine how reactor-array geometry alters the effects of multilevel selection and how this, in turn, influences the fate of polymer ecosystems.

In this study, we only considered two types of polar monomers that match in a reverse-complementary manner. However, real RNAs use four bases, raising polymer diversity at all lengths. It would also be worth exploring alternative potential mechanisms of template-guided replication, if only to evaluate whether reverse-complementary base-pairing is uniquely suited to the emergence of genetic encoding. For example, one could readily consider non-RNA-based models in which monomers and polymers are nonpolar, models in which base-pairing is directly complementary (e.g., ABB matching BAA instead of AAB), or even models where pairing happens between monomers of the same type (e.g., A matching A, B matching B), etc. To simulate these alternative scenarios, one could simply make changes to the adsorption phase and the rules determining reactive borders (Fig. 1).

We only considered three catalytic functions in this work: kinase, ligase, and mutation inhibition (which corrects wildcards), but many more biochemical functions are conceivable. For example, one may modify the reaction phase (Fig. 1) to make the synthesis of free activated monomers consist of more steps, better mimicking complex metabolic pathways. Similarly, we could allow catalysts to bind to substrates so as to allow phenomena such as processive polymerization, although doing this may require a more complicated module tracking the status of adsorbed molecules. Alternatively, we could allow polymers to confer structural rather than catalytic functions, for example changing surface properties in such a way as to reduce the rate of loss by dilution.

The approach taken here, of seeding ecosystems with catalytic polymers, is sufficient to explore diverse factors influencing catalyst persistence. However, if we want to understand how new functional sequences could arise over time, we will need to modify the simulation engine developed here to simulate open-ended adaptive evolution. A key addition would be a module automatically assigning catalytic functions (of diverse kinds) and catalytic efficiencies to newly emerged sequences, in such a way that longer sequences have higher maximum catalytic efficiencies. Using such a module, one could potentially study the process by which the first catalysts arise and are later replaced by new, more efficient (and perhaps longer) polymers. Such an approach should make it possible to explore long-term complexification in ecosystems of cooperating polymers, all driven by multilevel selection acting via reactor/ecosystem productivity. Such an extension would greatly advance our understanding of the origins of life by better explaining the co-emergence and co-complexification of metabolisms and genetic information processing systems.

## Supporting information

Supplemental Materials, Methods, Table & References

Supplemental Python Scripts

## Acknowledgement

We thank the Center of High Throughput Computing at the University of Wisconsin–Madison for assistance in parallel computing. We also thank Emily Dolson, Eric Smith, and Praful Gagrani for helpful discussions. This study was supported by the National Science Foundation (DEB-2218817) and the National Aeronautics and Space Administration (80NSSC17K0296).

