## Supplemental Materials, Methods, Table & References for "Conditions enabling the persistence of cooperating ribozymes without cellular encapsulation"


### Authors & Affiliations

Zhen Peng[1], Alex M. Plum[1,2,†], Rahul Kartha[1,3,‡], Emily M. Jacobson[1,2], David A. Baum[1,4]*

[1]Wisconsin Institute for Discovery, University of Wisconsin–Madison, Madison, WI 53706, USA

[2]Department of Engineering Physics, University of Wisconsin–Madison, Madison, WI 53706, USA

[3]Department of Computer Sciences, University of Wisconsin–Madison, Madison, WI 53706, USA

[4]Department of Botany, University of Wisconsin–Madison, Madison, WI 53706, USA

[†]Current affiliation: Department of Physics, University of California San Diego, La Jolla, CA 92093, USA

[‡]Current affiliation: Meta Platforms, Inc., 1 Meta Way, Menlo Park, CA 94025, USA

**Supplemental Materials & Methods**

The model simulates reactions in one or multiple reactors across multiple generations. Between two generations, partial replacement of matter in a reactor by external food source and inter-reactor exchange of matter may occur. Within a generation, multiple three-phase cycles are iterated to synthesize activated monomers from foods and new polymers from activated monomers.

In this model, except for the diverse polymers, eight chemical species are considered: energetic food E, structural foods F and G, activated monomers A and B, activated wildcards M and N, and waste W. Of these, all except energetic food and waste are considered polar, meaning that when adsorbed, reactivity differs between the two adjacent sites.

Ignoring ligation and polymer degradation reactions, only three classes of reactions are allowed:

1. Activation and deactivation of monomers: $E + F \rightleftharpoons A + W$, $E + G \rightleftharpoons B + W$,
2. Introduction of wildcards: $A \rightarrow M$, $B \rightarrow N$ (including the monomers incorporated in polymers),
3. Correction of wildcards: $M \rightarrow A$, $N \rightarrow B$ (including the monomers incorporated in polymers).

**1. Within a generation**

In all simulations reported in the paper, the number of cycles through the following phases was set to 300.

_Phase 1: Adsorption (including production and correction of wildcards)_

*Stage 1a: Production of wildcards*

A and B have a fixed probability $P_{\text{int}}$, which is the wildcard-introduction probability, to become wildcards M and N, respectively. For the results reported in Section 2.6, $P_{\text{int}}$ varies as specified; for other simulations, $P_{\text{int}} = 10^{-6}$.

*Stage 1b: Correction of wildcards (extended model only)*

In a reactor, assuming that there is a polymer species capable of inhibiting mutation by converting a wildcard back to a normal monomer, using $x$ to denote the count of the mutation inhibitors and using $P_{\text{keep}}$ to denote the probability that a wildcard is **NOT** converted back to a normal monomer, inhibitors are taken to have the following effects on a wildcard:

1) If there is no inhibitor, the wildcard will never be converted back, which means $P_{\text{keep}}(x=0) = 1$;

2) More inhibitors (i.e., larger $x$) are associated with higher probability that the wildcard is converted back (i.e., smaller $P_{\text{keep}}$), which means $\Delta P_{\text{keep}}/\Delta x < 0$;

3) If there are unlimited inhibitors, the wildcard will surely be converted back, which means $P_{\text{keep}}(x \rightarrow \infty) = 0$.

Any function $P_{\text{keep}}(x)$ that satisfies these three conditions could be used, each representing a different way to scale the same underlying $P_{\text{keep}} \sim x$ relationship. Here we used a simple function:

$$P_{\text{keep}}(x) = (1 - v)^x, 0 < v < 1 \tag{1}$$

where $v$ is an index of the efficiency of wildcard correction, with higher $v$ resulting in a lower realized mutation rate.

Although in this paper, we only considered one mutation inhibitor species, it is not hard to derive a model that would allow for multiple mutation inhibitor species. If there are $H$ polymer species capable of inhibiting mismatch-induced mutation and inhibitors act independently, then

$$P_{\text{keep}}(x_1, x_1, \dots, x_i, \dots, x_H) = \prod_{i=1}^{H} (1 - v_i)^{x_i}, 0 < v_i < 1 \tag{2}$$

Then the probability $P_{\text{wild}}$ that a monomer remains a wildcard can be estimated by

$$P_{\text{wild}} = P_{\text{int}} \cdot P_{\text{keep}}(x_1, x_1, \dots, x_i, \dots, x_H) = P_{\text{int}} \cdot \prod_{i=1}^{H} (1 - v_i)^{x_i}, 0 < v_i < 1 \tag{3}$$

Also note that under such implementation, a mutation inhibitor is reusable.

*Stage 1c: Adsorption, with match and mismatch*

A reactor is assumed to have a very narrow bottom surface separated into $m$ sites. For Table S1, $m = 200$; for simulations with 30-reactor arrays, $m = 100$; for simulations with only one giant reactor, $m = 3000$.

Each site may adsorb a monomer A, B, M, or N (wheter free or incorporated in a polymer), a food molecule E, F, or G, or a waste molecule W to form first-layer adsorbates. All polymers and their monomers are polar. It is also assumed that the total count of nonpolar species is always more than $m$, which means that first-layer adsorption will always be saturated.

It is assumed that once a polar species is adsorbed to a site as a first-layer adsorbate, it has a probability $P_{\text{dead}}$ to convert an adjacent unoccupied site into a "dead" site that cannot adsorb other polar species (i.e., a "dead" site is accessible only to E and W, which are nonpolar). A lower $P_{\text{dead}}$ means that the free ends of first-layer monomers are more likely to be adjacent, which leads to a high probability of fusion between second-layer reverse complements that match first-layer templates, resulting in fusion-induced mutation. We implemented the appearance of "dead" sites in order to control the frequency of fusion-induced mutation. One possible physical interpretation for such "dead" sites is that polar molecules could adopt some tautomers or conformational isomers that sterically modify the properties of adjacent sites. When illustrating the ability of the model to handle long polymers (Table S1), we let $P_{\text{dead}} = 10^{-6}$; for other simulations, we let $P_{\text{dead}} = 1 - 10^{-6}$.

To simulate first-layer adsorption, **first** an unadsorbed molecule is randomly picked and randomly put at a site of the reactor. If the molecule is not a polymer, it will occupy only that site; otherwise, it will occupy multiple sites. For polar molecules there is a 50:50 chance of being placed in either orientation. **Second**, if the newly adsorbed molecule is polar, whether a "dead" adjacent site is created will be stochastically determined according to $P_{\text{dead}}$; if it is created, then the "dead" site will only accept nonpolar species for the remainder of the cycle; otherwise, the adjacent site is also accessible to polar species. **Third**, according to the results of the second step, new molecules are randomly picked and put on the sites adjacent to the sites that are already occupied. One may imagine this process as that the first adsorbate serves as a nucleate core, and the following adsorbates gradually expand their occupancy until the surface is saturated. **Fourth,** the second and third steps are iterated until all first-layer sites are occupied.

After first-layer adsorption ends, second-layer adsorption starts. Since second-layer adsorption depends on reverse complementary matching between monomers, only polymers and free monomers can be in the second layer. And for clarity, we will use "x" and "y" flanking A or B to mark the orientation of a monomer, and "v" to mark a free end of a monomer; for example, vxAyv is a free monomer A, and vxAyxByv is a free dimer AB. **First**, all possible matches between first-layer templates and monomers/polymers that are not yet adsorbed (i.e., candidate second-layer adsorbates) are computed and stored in a list that will be referred to as the matching list. In this list, each match is associated with a weight that is the count of the candidate second-layer adsorbate in the match. For example, assuming that the first layer only has 100 sites and they are completely occupied by one polymer consisting of 100 A's (denoted by $A_1A_2 \ldots A_{100}$ for clarity), and that there are 10 BBB's and 20 BBBB's waiting to be adsorbed, then there are 98 possible matches for a BBB (e.g., BBB matching $A_{98}A_{99}A_{100}$), each associated with the weight 10, and there are 97 possible matches for a BBBB, each associated with the weight 20. The matching follows these rules: (1) Every monomer in the second layer must match a monomer (not food, waste, or dead sites) in the first layer. (2) The matching requires correct orientation. For example, if two adjacent first-layer sites are occupied by monomers with different orientation [vxAyv, vyBxv], then no polymer can match these two monomers; alternatively, if the first-layer monomers have the same orientation [vxAyv, vxByv], then the dimer vxAyxByv (i.e., vyBxyAxv) can match them but the dimer vxByxAyv (i.e., vyAxyBxv) cannot. (3) The allowed matching pairs include A-B, A-M, A-N, B-M, B-N, M-M, M-N, and N-N. **Second**, one match is randomly picked from the matching list proportional to its weight. **Third**, the picked match is implemented. **Fourth**, the match and any other matches that it blocks are removed from the list (i.e., the picked match occupies second-layer sites that are also required by some other candidate matches) and all remaining matches that involve the newly adsorbed species have their weights decreased by 1. **Fifth**, the process continues until no further matches are possible.

After second-layer adsorption ends, the molecules that are still not adsorbed are assumed to be in the aqueous phase, moving around and colliding with the first-layer and second-layer borders between sites. And all M and N are converted back to A and B, respectively.

*Phase 2: Reaction*

There are two types of reversible reactions: (I) the production of an activated monomer and waste from an energetic food and a structural food, and its reverse, (II) the formation of a bond between monomers, and its reverse. Type (I) reactions only happen in the first layer and both reactants are required to be exposed to the aqueous phase, whereas Type (II) reactions only happen in the second layer since the first-layer monomers and polymers have their backbones protected by the bottom surface.

For clarity, we will use "z" to mark the orientation of a polar food.

**First**, all borders between sites are screened to find reactive borders according to the following rules. **(1)** If a first-layer border is between a nonpolar molecule (i.e., E or W) and a polar food (i.e., zF or zG) or a free monomer (i.e., vxAyv or vxByv), and the z-side or vx-side faces the nonpolar molecule, then the border is reactive; note that although W and zF/zG do not react and E and vxAyv/vxByv do not react, if the W or E happens to react with a molecule on the other side and becomes E or W, respectively, then the next reaction may happen to the focal border. **(2)** If a second-layer border is between monomers of the same orientation, then the border is reactive. Then the positions of the two types of reactive borders are stored in two lists.

**Second**, the total number of collisions between catalyst species (both environmental catalysts and catalytic polymers) and borders (whether they are reactive or not) actually used in the current cycle, $N_{\text{act.total}}$, is estimated according to

$$N_{\text{act.total}} \sim \begin{cases} \text{Poisson}(\lambda = N_{\text{total}}), \lambda \leq 10^9 \\ \lfloor \text{Normal}(\mu = N_{\text{total}}, \sigma^2 = N_{\text{total}}) \rfloor, \mu > 10^9 \end{cases} \tag{4}$$

where $N_{\text{total}}$ is the average total number of collisions. And $N_{\text{total}}$ is estimated according to

$$N_{\text{total}} = c_{\text{env.kin.}} \cdot L_{\text{surface}} + c_{\text{env.lig.}} \cdot L_{\text{surface}} + \sum_i c_{\text{base}} \cdot \frac{x_i}{\sqrt{y_i}} \tag{5}$$

where $L_{\text{surface}}$ is the length of the bottom surface, $c_{\text{env.kin.}}$ and $c_{\text{env.lig.}}$ are constants representing the joint effect of the concentration and diffusivity of environmental catalysts ("natural kinase" and "natural ligase") that catalyze spontaneous reversible activation and ligation reactions, respectively, $c_{\text{base}}$ is a constant used to convert the physical properties of catalytic polymers to collision numbers, $x_i$ is the count of the catalytic polymer species $i$, and $y_i$ is the number of monomers in the catalytic polymer species $i$.

The derivation of Equations (4) and (5) can be found in "**3. Derivation of Equations (4) and (5)**" of Supplemental Materials & Methods.

**Third**, $N_{\text{act.total}}$ collision events are simulated as follows. **(1)** The catalyst species colliding with borders between sites is randomly picked according to

$$p_{\text{env.kin.}} = \frac{c_{\text{env.kin.}} \cdot L_{\text{surface}}}{N_{\text{total}}} \tag{6}$$

$$p_{\text{env.lig.}} = \frac{c_{\text{env.lig.}} \cdot L_{\text{surface}}}{N_{\text{total}}} \tag{7}$$

$$p_i = \frac{c_{\text{base}}}{N_{\text{total}}} \cdot \frac{x_i}{\sqrt{y_i}} \tag{8}$$

where $p_{\text{env.kin.}}$, $p_{\text{env.lig.}}$, and $p_{i,y}$ are the probabilities of picking "natural kinase", "natural ligase", and catalytic polymer species *i*, respectively. **(2)** Once a catalyst species is picked, a border will be randomly picked (no matter it is reactive or not); if for the picked catalyst, the picked border happens to be reactive (e.g., first-layer [E, zF/zG] and [W, vxAyv/vxByv] for kinase, second-layer [yAx, yBxv] and [yAxv, vyBxv] for ligase), proceed to the next step; otherwise, go back to the previous step. **(3)** The catalyst has an intrinsic probability to make the reaction occur, and whether the reaction actually occurs is stochastic; such a probability is considered as the catalytic efficiency in this study. Since all reactions are assumed to be reversible, every catalyst has a pair of such efficiencies; and for the same type of catalysts, their ratios of forward efficiency to reverse efficiency keep constant due to thermodynamics (i.e., catalysts do not modify the change in the Gibbs energy); for kinase, the ratio of activation to deactivation is 1:1; for ligase, the ratio of ligation to breakage is $10^4$:1. For Table S1, the efficiencies of spontaneous activation and spontaneous ligation are both 0.9; for other simulations, they are both $10^{-5}$. If the reaction does not occur, then go back to step (1); however, if the reaction occurs, then the affected adsorbates will be updated before going back to step (1): activation converts [E, zF/zG] to [W, vxAyv/vxByv], while deactivation does the reverse; ligation removes "v"s flanking the border, while breakage does the reverse. **(4)** The previous three steps are iterated until all $N_{\text{act.total}}$ collision events occur.

*Phase 3: Resuspension*

All adsorbates are released to the aqueous phase.

**2. Between generations**

Once the resuspension phase of the last cycle in a generation is completed, every reactor will have a proportion of its molecules stochastically lost, and the same proportion of molecules from an external source container with the same volume stochastically added to the reactor. This proportion is the dilution rate. For example, assuming a dilution rate of 0.1, if a unit-volume reactor has 800 molecules and a unit-volume external source has 1000 E, 500 F, and 500 G, then it is expected that 80 old molecules will be lost while 100 E, 50 F, and 50 G will enter the reactor.

After dilution, reactors will enter the period when inter-reactor exchange of matter may occur, with three possible regimes.

*Regime 1: Neighborhood dispersal*

Molecules stochastically move to adjacent reactors or stay in the local reactor. In this study, we assumed that the probability of staying is 0.6, the probability of moving to the left reactor is 0.2, and the probability of moving to the right reactor is also 0.2. The reactor array is assumed to be circular, meaning that the reactor to the left of the leftmost reactor is the rightmost reactor.

*Regime 2: Global dispersal*

Molecules stochastically stay or enter a shared pool; in the shared pool, molecules from different reactors are well-mixed and then stochastically distributed to all reactors. In this study, the probability of staying is 0.6 while probability of entering the shared pool is 0.4.

*Regime 3: Well-mixed giant reactor*

Since there is only one reactor, inter-reactor exchange does not occur.

**3. Derivation of Equations (4) and (5)**

Since we assume that all reactors have the same height, according to Fick's law of diffusion, the number of collisions between Brownian particles and the bottom surface of the reactor per unit area per unit time, $F_{\text{collision}}$, should follow

$$F_{\text{collision}} \propto D \cdot \frac{x}{V} \tag{9}$$

where $D$ is the diffusivity, $x$ is the count of particles, and $V$ is the volume of the container. Therefore, we get the following estimation

$$F_{\text{collision}} = D \cdot \frac{x}{V} \cdot c_1 \tag{10}$$

where $c_1$ is a constant and its unit is $m^{-1}$. Since different reactors are rectangular cuboids that share the same width and the same height, we have

$$L_{\text{surface}} \propto A_{\text{surface}} \propto V \tag{11}$$

where $L_{\text{surface}}$ is the length of the bottom surface, and $A_{\text{surface}}$ is the area of the bottom surface. Then $F_{\text{collision}}$ can be estimated by

$$F_{\text{collision}} = c_1 c_2 \cdot \frac{x}{A_{\text{surface}}} D \tag{12}$$

where $c_2$ is a constant and its unit is $m^{-1}$. Then the number of collisions between particles and the bottom surface per particle per unit time, $r_{\text{collision.surf}}$, can be estimated by

$$r_{\text{collision.surface}} = F_{\text{collision}} \cdot A_{\text{surface}} \cdot \frac{1}{x} = c_1 c_2 \cdot \frac{x}{A_{\text{surface}}} D \cdot A_{\text{surface}} \cdot \frac{1}{x} = c_1 c_2 D \tag{13}$$

Denoting the number of collisions between particles and borders between sites per particle per unit time by $r_{\text{collision}}$, and the number of collisions between particles and sites per particle per unit time by $r_{\text{collision.site}}$, we have

$$r_{\text{collision.surface}} = r_{\text{collision}} + r_{\text{collision.site}} \tag{14}$$

Then denoting the number of sites on the bottom surface by $m$, the number of borders between sites by $b$, the surface area of a site by $a_m$ which is a constant, and the surface area of a border by $a_b$ which is a constant (note that the borders between sites have some finite size), we have

$$\frac{r_{\text{collision}}}{r_{\text{collision.site}}} = \frac{b a_b}{m a_m} = \frac{m-1}{m} \cdot \frac{a_b}{a_m} = \frac{m-1}{m} \cdot \frac{1}{c_{mb}} \tag{15}$$

where $c_{mb}$ is a constant. According to Equations (14) and (15), we get

$$r_{\text{collision}} = \frac{r_{\text{collision.surface}}}{1 + \frac{m}{m-1} \cdot c_{mb}} = \frac{c_1 c_2 D}{1 + \frac{m}{m-1} \cdot c_{mb}} \tag{16}$$

In our simulations, the minimum $m$ is 100, which is large enough to accept the approximation

$$\frac{m}{m-1} \approx 1 \tag{17}$$

Consequently, we get the following estimation

$$r_{\text{collision}} = \frac{c_1 c_2 D}{1 + c_{mb}} = c_3 \cdot c_1 c_2 D \tag{18}$$

where $c_3$ is a constant. Now assuming that the temperature keeps constant, it is obvious that $D$ can be considered as a constant for a given chemical species, and thus $r_{\text{collision}}$ is a constant.

Now consider two chemical species $i$, $j$ that are in the same container but have different lengths $l_i$, $l_j$. According to Equation (18), we have

$$\frac{r_{\text{collision},i}}{r_{\text{collision},j}} = \frac{c_1 c_2 c_3 D_i}{c_1 c_2 c_3 D_j} = \frac{D_i}{D_j} \tag{19}$$

The relationship between a polymer's diffusivity and its length is usually considered to satisfy

$$D \propto l^{-\nu} \tag{20}$$

where $\nu$ is a positive constant (Robertson et al., 2006). Depending on specific assumptions, such as how a polymer chain folds into a particle, $\nu$ will take different values. For simplification, here we adopt the ideal chain model in which $\nu = 1/2$ (de Gennes, 1979, pp. 29–42), then we get

$$\frac{r_{\text{collision},i}}{r_{\text{collision},j}} = \frac{D_i}{D_j} = \left(\frac{l_i}{l_j}\right)^{-1/2} = \sqrt{\frac{l_j}{l_i}} \tag{21}$$

Assuming that for a free monomer, its number of collisions with borders per particle per unit time is a constant $r_{\text{collision},1}$, we can estimate that for a $y$-mer species $i$, its $r_{\text{collision},i,y}$ is

$$r_{\text{collision},i,y} = r_{\text{collision},1} \sqrt{\frac{l_1}{l_{i,y}}} = r_{\text{collision},1} \sqrt{\frac{1}{y}} = \frac{r_{\text{collision},1}}{\sqrt{y}} \tag{22}$$

Then the total number of collisions between a $y$-mer species $i$ and borders during the entire reaction phase, $N_{i,y}$, is given by

$$N_{i,y} = r_{\text{collision},i,y} \cdot t \cdot x_i = r_{\text{collision},1} \cdot t \cdot \frac{x_i}{\sqrt{y_i}} \tag{23}$$

where $t$ is the timespan of the reaction phase, which is a constant. Then let

$$c_{\text{base}} = r_{\text{collision},1} \cdot t \tag{24}$$

Then we can use the approximation

$$N_{i,y} = c_{\text{base}} \cdot \frac{x_i}{\sqrt{y_i}} \tag{25}$$

For all catalytic polymers, their $N_{i,y}$'s are computed. In this study, we set $c_{\text{base}} = 10^4$.

To implement spontaneous reactions, we assume that there exist readily available environmental "kinases" and "ligases" that have fixed concentrations in the food source, then it is easy to derive the approximation

$$N_{\text{env.kin.}} = c_{\text{env.kin.}} \cdot L_{\text{surface}} \tag{26}$$

$$N_{\text{env.lig.}} = c_{\text{env.lig.}} \cdot L_{\text{surface}} \tag{27}$$

where $N_{\text{env.kin.}}$ is the total number of collisions between environmental kinases and borders, $N_{\text{env.lig.}}$ is the total number of collisions between environmental ligases and borders, and $c_{\text{env.kin.}}$ and $c_{\text{env.lig.}}$ are constants. For Table S1, we set $c_{\text{env.kin.}} = c_{\text{env.lig.}} = 5\times10^4$ (per unit length); for other simulations, we set $c_{\text{env.kin.}} = c_{\text{env.lig.}} = 1$ (per unit length); here, a unit length equals the sum of 100 sites.

Next, the total number of collisions between catalysts and borders, $N_{\text{total}}$, is computed by

$$N_{\text{total}} = N_{\text{env.kin.}} + N_{\text{env.lig.}} + \sum_i N_{i,y} \tag{28}$$

Substituting the corresponding terms in Equation (28) by Equations (25), (26), and (27), we get

$$N_{\text{total}} = c_{\text{env.kin.}} \cdot L_{\text{surface}} + c_{\text{env.lig.}} \cdot L_{\text{surface}} + \sum_i c_{\text{base}} \cdot \frac{x_i}{\sqrt{y_i}} \tag{5}$$

When performing stochastic simulation, $N_{total}$ is treated as the expected number of collisions rather than the actual total number of collisions. Since we care about how many events (i.e., collisions) happen within a time interval (i.e., one reaction phase), we can use Poisson distribution to estimate the actual total number of collisions between catalysts and borders, $N_{act.total}$; and if $N_{total}$ is large enough, a normal distribution can also be used. Consequently, we get

$$N_{\text{act.total}} \sim \begin{cases} \text{Poisson}(\lambda = N_{\text{total}}), \lambda \leq 10^9 \\ \lfloor \text{Normal}(\mu = N_{\text{total}}, \sigma^2 = N_{\text{total}}) \rfloor, \mu > 10^9 \end{cases} \quad (4)$$

**Supplemental Table**

| Replicate | Mean Length | Median Length | Maximum Length |
|---|---|---|---|
| 1 | 21.051 | 19 | 54 |
| 2 | 17.382 | 16 | 34 |
| 3 | 19.642 | 20 | 63 |
| 4 | 21.837 | 20 | 57 |
| 5 | 18.927 | 17 | 36 |
| 6 | 18.796 | 17 | 39 |
| 7 | 17.125 | 17 | 47 |
| 8 | 20.079 | 19.5 | 56 |
| 9 | 17.250 | 16.5 | 32 |
| 10 | 21.976 | 21 | 47 |
| 11 | 18.769 | 16 | 33 |
| 12 | 18.525 | 16 | 56 |
| 13 | 19.118 | 18 | 41 |
| 14 | 18.232 | 17 | 41 |
| 15 | 19.360 | 18 | 39 |
| 16 | 21.451 | 19 | 50 |
| 17 | 20.042 | 18 | 48 |
| 18 | 21.646 | 20.5 | 47 |
| 19 | 16.417 | 15 | 33 |
| 20 | 18.833 | 17 | 51 |
| 21 | 18.717 | 18 | 45 |
| 22 | 18.327 | 16 | 36 |
| 23 | 20.302 | 19 | 46 |
| 24 | 17.407 | 17 | 39 |
| 25 | 20.229 | 18.5 | 50 |

| | | | |
|---|---|---|---|
| 26 | 19.286 | 15 | 45 |
| 27 | 20.923 | 18 | 49 |
| 28 | 17.533 | 16 | 39 |
| 29 | 19.143 | 18 | 51 |
| 30 | 19.528 | 19 | 35 |
| Average | 19.262 | 17.733 | 44.633 |
| Standard Deviation | 1.488 | 1.596 | 8.227 |

**Table S1. Demonstration that the basic model can handle long polymers.** Each replicate has one reactor with 200 sites. Each replicate is initialized by 1000 E, 500 F, and 500 G. Simulations ran for 20 generations; inter-generation dilution rate was 0.05; the rates of spontaneous reactions were set to high levels ($c_{\text{env.kin.}} = c_{\text{env.lig.}} = 5\times10^4$ (per unit length), $v_{\text{env.kin.}} = v_{\text{env.lig.}} = 0.9$); the probability of fusion-induced mutations was set to high levels (i.e., the probability of creating "dead" sites is allow, $P_{\text{dead}} = 10^{-6}$).
