## Supplemental Python Scripts for "Conditions enabling the persistence of cooperating ribozymes without cellular encapsulation": Merge_Distri.sh

```
#!/bin/bash

# untar your Python installation. Make sure you are using the right version!
#tar -xzf python38.tar.gz
# (optional) if you have a set of packages (created in Part 1), untar them also
#tar -xzf np.tar.gz

# make sure the script will use your Python installation, 
# and the working directory as its home location
export PATH=$PWD/python/bin:$PATH
export PYTHONPATH=$PWD/packages
export HOME=$PWD

# run your script
python3 $1 $2 $3 $4 $5

#python3 Merge_Distri.py Num_reactors Replacement_ratio Current_generation
```
