## Supplemental Python Scripts for "Conditions enabling the persistence of cooperating ribozymes without cellular encapsulation": Merge_Distri_period_neighbor.py

```
'''
python Merge_Distri.py reactor_num permanent_loss_ratio local_stay_ratio generation_num 

test:
python Merge_Distri.py 50 0.5 0.6 1 > 00_testout.txt

'''
import glob
#import os
import sys
#import time
import numpy as np


volume = 1


food_profile = {
'E':1000*volume,
'F':500*volume,
'G':500*volume,
}

reactor_num = int(sys.argv[1])
permanent_loss_ratio = float(sys.argv[2])
local_stay_ratio = float(sys.argv[3])
generation_num = int(sys.argv[4])
#output_dir = sys.argv[4]


def convert_list_to_count_dict(ob_list):
	count_dict = {}
	for ob in ob_list:
		if ob not in count_dict:
			count_dict[ob] = 1
		else:
			count_dict[ob] = count_dict[ob] + 1
	return(count_dict)


def distribute_focal_reactor_mol_to_next_generation_reactors(next_generation_reactors_array, current_reactor_mol_list, focal_reactor_index, permanent_loss_ratio, local_stay_ratio):
	reactor_num = len(next_generation_reactors_array)
	prob_loss = permanent_loss_ratio
	prob_stay = (1-permanent_loss_ratio)*local_stay_ratio
	prob_move_left = (1 - prob_loss - prob_stay)/2

	for mol in current_reactor_mol_list:
		fate = np.random.choice(['Loss','Stay','Left','Right'], p=[prob_loss, prob_stay, prob_move_left, 1-prob_loss-prob_stay-prob_move_left])
		if fate=='Stay':
			target_reactor_index = focal_reactor_index
			if mol in next_generation_reactors_array[target_reactor_index]:
				next_generation_reactors_array[target_reactor_index][mol] = next_generation_reactors_array[target_reactor_index][mol]+1
			else:
				next_generation_reactors_array[target_reactor_index][mol] = 1
		elif fate=='Left':
			#if focal_reactor_index>0:
			target_reactor_index = focal_reactor_index-1
			if mol in next_generation_reactors_array[target_reactor_index]:
				next_generation_reactors_array[target_reactor_index][mol] = next_generation_reactors_array[target_reactor_index][mol]+1
			else:
				next_generation_reactors_array[target_reactor_index][mol] = 1
		elif fate=='Right':
			if focal_reactor_index<reactor_num-1:
				target_reactor_index = focal_reactor_index+1
			else:
				target_reactor_index = 0
			
			if mol in next_generation_reactors_array[target_reactor_index]:
				next_generation_reactors_array[target_reactor_index][mol] = next_generation_reactors_array[target_reactor_index][mol]+1
			else:
				next_generation_reactors_array[target_reactor_index][mol] = 1

	return(next_generation_reactors_array)


def distribute_food_to_reactors(next_generation_reactors_array, permanent_loss_ratio, local_stay_ratio, food_profile):
	food_mol_list = []
	for food in food_profile:
		food_raw = food_profile[food]
		for m in range(food_raw):
			food_mol_list.append(food)
	food_mol_num = len(food_mol_list)
	
	reactor_num = len(next_generation_reactors_array)
	mid_reactor_food_enter_prob = permanent_loss_ratio
	#edge_reactor_food_enter_prob = permanent_loss_ratio + (1-permanent_loss_ratio)*(1-local_stay_ratio)/2

	for i in range(reactor_num):
		#if i in [0, reactor_num-1]:
		#	for food_name in food_mol_list:
		#		fate = np.random.choice(['Enter','Miss'], p=[edge_reactor_food_enter_prob, 1-edge_reactor_food_enter_prob])
		#		if fate=='Enter':
		#			if food_name in next_generation_reactors_array[i]:
		#				next_generation_reactors_array[i][food_name] = next_generation_reactors_array[i][food_name]+1
		#			else:
		#				next_generation_reactors_array[i][food_name] = 1
		#else:
		for food_name in food_mol_list:
			fate = np.random.choice(['Enter','Miss'], p=[mid_reactor_food_enter_prob, 1-mid_reactor_food_enter_prob])
			if fate=='Enter':
				if food_name in next_generation_reactors_array[i]:
					next_generation_reactors_array[i][food_name] = next_generation_reactors_array[i][food_name]+1
				else:
					next_generation_reactors_array[i][food_name] = 1

	return(next_generation_reactors_array)


next_generation_reactors_array = []
for k in range(reactor_num):
	next_generation_reactors_array.append({})

next_generation_reactors_array = distribute_food_to_reactors(next_generation_reactors_array, permanent_loss_ratio, local_stay_ratio, food_profile)


last_line_dict = {}

for g in glob.glob('*.out'):
	if "SIMULATION" in g:
		focal_reactor_name = g.split('.')[0]
		focal_reactor_name = focal_reactor_name.split('_')[1]
		focal_reactor_index = int(focal_reactor_name)
		
		current_reactor_mol_list = []
		#print(g)
		last_line = ''
		outfile = open(g,'r')
		for line in outfile:
			line = line.strip()
			if len(line)>0:
				last_line = line
		outfile.close()

		last_line_dict[focal_reactor_index] = last_line

		record_split = last_line.split(', ')
		for name_count in record_split:
			[name,count] = name_count.split(':')
			count = int(count)
			for i in range(count):
				current_reactor_mol_list.append(name)

		next_generation_reactors_array = distribute_focal_reactor_mol_to_next_generation_reactors(next_generation_reactors_array, current_reactor_mol_list, focal_reactor_index, permanent_loss_ratio, local_stay_ratio)


for reactor_index in range(reactor_num):
	
	mol_dict = next_generation_reactors_array[reactor_index]

	mol_name_list = []
	for mol in mol_dict:
		mol_name_list.append(mol)
	mol_name_list.sort()
	mol_name_list = sorted(mol_name_list, key=len)

	print_list = []
	for mol in mol_name_list:
		mol_count = mol_dict[mol]
		print_ob = mol+':'+str(mol_count)
		print_list.append(print_ob)
	print_line = ', '.join(print_list)

	#file_name = 'Generation_'+str(generation_num+1)+'/distri_'+reactor_index+'.txt'
	#os.makedirs(os.path.dirname(file_name), exist_ok=True)
	#file = open(file_name,'w')
	print(print_line)
	#file.close()

print()
print('#=========== Last_Generation_Final_Counts ===========#')
print()
for reactor_index in range(reactor_num):
	print(last_line_dict[reactor_index])

#file_name = 'distri_'+str(i)+'.txt'

#np.random.shuffle(food_mol_list)
#total_food_num = len(food_mol_list)
#food_num_enter_next_gen = int(food_mol_list*(1-replacement_ratio))
#food_enter_next_gen_list = molecule_list[0:mol_num_enter_next_gen]
```
