## Supplemental Python Scripts for "Conditions enabling the persistence of cooperating ribozymes without cellular encapsulation": Simu_Tetris_one_reactor_9optm.py

```
'''
python Simu_Tetris_one_reactor_ParamTest.py distri.txt reactor_index > Tetris_one_reactor_out1.txt

Test:
python Simu_Tetris_one_reactor_ParamTest.py distri.txt 0 > Tetris_one_reactor_out1.txt

1-D surface for simplification

This script only applies to cases where the count of molecules is much more than the surface length


python Simu_Tetris_one_reactor_9optm.py test_distri.txt 0 > out_1stLayer.txt


python Simu_Tetris_one_reactor_9optm.py test_distri_X1.txt 0 > out_1stLayer.txt


python Simu_Tetris_one_reactor_9optm.py test_distri_X2.txt 0 > out_1stLayer.txt

python Simu_Tetris_one_reactor_9optm.py test_distri_X3.txt 0 > out_WlmxTest1.txt

python Simu_Tetris_one_reactor_9optm.py test_distri_X4.txt 0 > out_WlmxTest2.txt

'''


import sys
import numpy as np


volume = 1
surface_length = 100*volume#80*volume#100#80
cycle_num = 300#250
base_collision_number_per_residue_per_cycle = 1e4#2500#1e3

# modify this to control the level of substitution-type mutations. 
mismatch_prob = 1e-6#0.1#1e-6 #0#1e-6#1e-4
# modify this to control the level of mergence-type mutations. 
D_site_prob = 1 - 1e-6#0.8#1 - 1e-6 #1#1 - 1e-6


kination_RF_ratio = 1
ligation_RF_ratio = 1e-4

# physical meaning: base_collision_number * concentration_of_natural_catalysts
natural_kination_catalyst_collisions_per_volume_per_cycle = 1
natural_ligation_catalyst_collisions_per_volume_per_cycle = 1

spontaneous_kination_prob = 1e-5
spontaneous_ligation_prob = 1e-5

kinase_dict = {
'ABAABABB':0.0005,
#'BABAABBBAA':0.005,
}

ligase_dict = {
'BBAAABAB':0.02,
#'BAABABAABB':0.2,
}


def convert_list_to_count_dict(ob_list):
	count_dict = {}
	for ob in ob_list:
		if ob not in count_dict:
			count_dict[ob] = 1
		else:
			count_dict[ob] = count_dict[ob] + 1
	return(count_dict)


def get_reverse_seq(seq):
	if len(seq)==1:
		reverse_seq = seq
	else:
		reverse_seq = ''
		seq_length = len(seq)

		#print('----------rev_mod-------------')
		#print(seq)
		#print('-----------check_end------------')


		for i in range(seq_length):
			reverse_seq = reverse_seq + seq[seq_length-i-1]
	return(reverse_seq)


def orient_polymer_seq(raw_seq):
	if raw_seq.startswith('vx'):
		xy_direction_seq = raw_seq
	else:
		xy_direction_seq = get_reverse_seq(raw_seq)
	return(xy_direction_seq)


def convert_normal_polymer_name_to_xy_name(normal_polymer_name):
	xy_name = 'v'+str(normal_polymer_name)+'v'
	xy_name = xy_name.replace('A','xAy')
	xy_name = xy_name.replace('B','xBy')
	return(xy_name)

def convert_xy_name_to_normal_polymer_name(xy_name):
	normal_polymer_name = orient_polymer_seq(xy_name)
	normal_polymer_name = normal_polymer_name.replace('v','')
	normal_polymer_name = normal_polymer_name.replace('x','')
	normal_polymer_name = normal_polymer_name.replace('y','')
	return(normal_polymer_name)


def inject_mismatch_residue(molecule_name_list, mismatch_prob):
	new_molecule_list = []
	for molecule in molecule_name_list:
		if 'x' not in molecule:
			new_molecule_list.append(molecule)
		else:
			molecule = orient_polymer_seq(molecule)
			tmp_str = ''
			for symbol in molecule:
				if symbol in ['A','B']:
					symbol = symbol + np.random.choice(['K',''], p = [mismatch_prob, 1-mismatch_prob])
				tmp_str = tmp_str+symbol
			tmp_str = tmp_str.replace('AK','M')
			tmp_str = tmp_str.replace('BK','N')
#=====================================
			#print(tmp_str)
#=====================================
			new_molecule_list.append(tmp_str)
	return(new_molecule_list)


def pick_first_molecule_from_molecule_count_dict(molecule_count_dict_with_injecting_mismatchable_residue, molecule_count):
	label_list = []
	p_list = []
	for label in molecule_count_dict_with_injecting_mismatchable_residue:
		prob = molecule_count_dict_with_injecting_mismatchable_residue[label]/molecule_count
		label_list.append(label)
		p_list.append(prob)
	
	#p_list[-1] = 1 - sum(p_list[0:-1])
	
	picked_molecule = np.random.choice(label_list, p = p_list)
	return(picked_molecule)


def pick_a_molecule_from_all_molecules(molecule_count_dict_with_injecting_mismatchable_residue, molecule_count):
	label_list = []
	p_list = []
	for label in molecule_count_dict_with_injecting_mismatchable_residue:
		prob = molecule_count_dict_with_injecting_mismatchable_residue[label]/molecule_count
		label_list.append(label)
		p_list.append(prob)
	
	#p_list[-1] = 1 - sum(p_list[0:-1])

	#print(p_list)

	picked_molecule = np.random.choice(label_list, p = p_list)
	return(picked_molecule)


def pick_a_molecule_from_EW(molecule_count_dict_with_injecting_mismatchable_residue):
	label_list = ['E','W']
	if 'E' in molecule_count_dict_with_injecting_mismatchable_residue:
		E_count = molecule_count_dict_with_injecting_mismatchable_residue['E']
	else:
		E_count = 0
	
	if 'W' in molecule_count_dict_with_injecting_mismatchable_residue:
		W_count = molecule_count_dict_with_injecting_mismatchable_residue['W']
	else:
		W_count = 0

	if E_count + W_count ==0:
		print('Error: No E or W!!!!!')
		sys.exit()

	p_E = E_count/(E_count + W_count)
	p_list = [p_E, 1-p_E]
	picked_molecule = np.random.choice(label_list, p = p_list)
	return(picked_molecule)


def get_molecule_length_from_name(molecule_name):
	molecule_length = 0
	for letter in molecule_name:
		if letter in ['A','B','E','F','G','W','M','N']:
			molecule_length = molecule_length + 1
	return(molecule_length)


def randomly_reverse_molecule(molecule):
	randnum = np.random.rand()
	if randnum<0.5:
		random_reversed_molecule = molecule
	else:
		random_reversed_molecule = get_reverse_seq(molecule)
	return(random_reversed_molecule)


def pick_following_molecule_from_molecule_count_dict(D_site_prob, current_molecule_name, molecule_count_dict_with_injecting_mismatchable_residue, molecule_count):
	#molecule_type = 'Undetermined'
	if 'x' in current_molecule_name:# monomer or polymer
		#molecule_type = 'polar'
		D_site_creation = np.random.choice(['Yes','No'], p=[D_site_prob, 1-D_site_prob])
		if D_site_creation=='No':# polar molecules are allowed, such as vxAyv, Fz, Gz
			next_molecule = pick_a_molecule_from_all_molecules(molecule_count_dict_with_injecting_mismatchable_residue, molecule_count)
		else:# nonpolar molecules only, E or W
			next_molecule = pick_a_molecule_from_EW(molecule_count_dict_with_injecting_mismatchable_residue)
	
	elif 'z' in current_molecule_name:# polar food Fz or Gz
		#molecule_type = 'polar'
		D_site_creation = np.random.choice(['Yes','No'], p=[D_site_prob, 1-D_site_prob])
		if D_site_creation=='No':# polar molecules are allowed, such as vxAyv, Fz, Gz
			next_molecule = pick_a_molecule_from_all_molecules(molecule_count_dict_with_injecting_mismatchable_residue, molecule_count)
		else:# nonpolar molecules only, E or W
			next_molecule = pick_a_molecule_from_EW(molecule_count_dict_with_injecting_mismatchable_residue)
	
	else:# E or W
		#molecule_type = 'nonpolar'
		next_molecule = pick_a_molecule_from_all_molecules(molecule_count_dict_with_injecting_mismatchable_residue, molecule_count)
	
	return(next_molecule)


def clean_tail_fill_in_gap(D_site_prob, occupied_length, surface_length, first_layer_molecule_list, molecule_count_dict_with_injecting_mismatchable_residue, molecule_count):
	#move the last molecule back to pool
	cleaved_molecule_label = first_layer_molecule_list[-1]
	first_layer_molecule_list = first_layer_molecule_list[:-1]	

	cleaved_molecule_length = get_molecule_length_from_name(cleaved_molecule_label)
	occupied_length = occupied_length - cleaved_molecule_length
	molecule_count_dict_with_injecting_mismatchable_residue[cleaved_molecule_label] = molecule_count_dict_with_injecting_mismatchable_residue[cleaved_molecule_label] + 1
	molecule_count = molecule_count + 1
	
	#pick and add molecules again
	current_molecule_name = first_layer_molecule_list[-1]

	#picker = 0
	while occupied_length<surface_length:
		next_molecule = pick_following_molecule_from_molecule_count_dict(D_site_prob, current_molecule_name, molecule_count_dict_with_injecting_mismatchable_residue, molecule_count)
		molecule_count_dict_with_injecting_mismatchable_residue[next_molecule] = molecule_count_dict_with_injecting_mismatchable_residue[next_molecule] - 1
		molecule_count = molecule_count - 1
		next_molecule_length = get_molecule_length_from_name(next_molecule)
		occupied_length = occupied_length + next_molecule_length
		#random_reversed_molecule = randomly_reverse_molecule(next_molecule)
		first_layer_molecule_list.append(next_molecule)
		current_molecule_name = next_molecule


	if occupied_length==surface_length:
		first_layer_ordered_molecules = first_layer_molecule_list
		#print('Good News! Exactly Matched!!!')
		#print('Occupied_length is '+str(occupied_length))
		#print(first_layer_molecule_list)


	elif occupied_length>surface_length:
#--------------------------------------------------------------------------------------
		#print('Indicator!!!: Tail cleaved and Gap filled!!!')
		#print('Occupied_length is '+str(occupied_length))
		#print(first_layer_molecule_list)
#--------------------------------------------------------------------------------------
		[first_layer_ordered_molecules, molecule_count_dict_with_injecting_mismatchable_residue, molecule_count] = clean_tail_fill_in_gap(D_site_prob, occupied_length, surface_length, first_layer_molecule_list, molecule_count_dict_with_injecting_mismatchable_residue, molecule_count)
	else:
		print("Error: Loop incomplete!!!")
		sys.exit() 

	return([first_layer_ordered_molecules, molecule_count_dict_with_injecting_mismatchable_residue, molecule_count])


def fill_in_first_layer_with_molecules(surface_length, D_site_prob, molecule_count_dict_with_injecting_mismatchable_residue, molecule_count):
	#occupied_length = 0
	first_layer_molecule_list = []
	first_molecule = pick_first_molecule_from_molecule_count_dict(molecule_count_dict_with_injecting_mismatchable_residue, molecule_count)
	molecule_count_dict_with_injecting_mismatchable_residue[first_molecule] = molecule_count_dict_with_injecting_mismatchable_residue[first_molecule] - 1
	molecule_count = molecule_count - 1
	first_molecule_length = get_molecule_length_from_name(first_molecule)
	#occupied_length = first_molecule_length
	#random_reversed_molecule = randomly_reverse_molecule(first_molecule)
	first_layer_molecule_list.append(first_molecule)

	current_molecule_name = first_molecule

	insert_site_index = np.random.choice(surface_length - first_molecule_length + 1)

	left_total_length = insert_site_index + first_molecule_length
	right_total_length = surface_length - insert_site_index


	# add molecules to the left
	left_occupied_length = first_molecule_length
	first_layer_left_half_molecule_list = [first_molecule]

	if left_total_length==left_occupied_length:
		first_layer_left_half_ordered_molecules = first_layer_left_half_molecule_list
	else:
		while left_occupied_length<left_total_length:
			next_molecule = pick_following_molecule_from_molecule_count_dict(D_site_prob, current_molecule_name, molecule_count_dict_with_injecting_mismatchable_residue, molecule_count)
			molecule_count_dict_with_injecting_mismatchable_residue[next_molecule] = molecule_count_dict_with_injecting_mismatchable_residue[next_molecule] - 1
			molecule_count = molecule_count - 1
			next_molecule_length = get_molecule_length_from_name(next_molecule)
			left_occupied_length = left_occupied_length + next_molecule_length

			first_layer_left_half_molecule_list.append(next_molecule)

			current_molecule_name = next_molecule

		if left_occupied_length==left_total_length:
			first_layer_left_half_ordered_molecules = first_layer_left_half_molecule_list
			#unused_molecule_count_dict_with_mismatchable_residue = molecule_count_dict_with_injecting_mismatchable_residue
		elif left_occupied_length>left_total_length:
			#[first_layer_ordered_molecules, unused_molecule_count_dict_with_mismatchable_residue] = clean_tail_fill_in_gap(D_site_prob, occupied_length, surface_length, first_layer_molecule_list, molecule_count_dict_with_injecting_mismatchable_residue, molecule_count)
			[first_layer_left_half_ordered_molecules, molecule_count_dict_with_injecting_mismatchable_residue, molecule_count] = clean_tail_fill_in_gap(D_site_prob, left_occupied_length, left_total_length, first_layer_left_half_molecule_list, molecule_count_dict_with_injecting_mismatchable_residue, molecule_count)

		else:
			print("Error: Loop incomplete!!!")
			sys.exit()


	# add molecules to the right
	right_occupied_length = first_molecule_length
	first_layer_right_half_molecule_list = [first_molecule]
	current_molecule_name = first_molecule
	if right_total_length==right_occupied_length:
		first_layer_right_half_ordered_molecules = first_layer_right_half_molecule_list
	else:
		while right_occupied_length<right_total_length:
			next_molecule = pick_following_molecule_from_molecule_count_dict(D_site_prob, current_molecule_name, molecule_count_dict_with_injecting_mismatchable_residue, molecule_count)
			molecule_count_dict_with_injecting_mismatchable_residue[next_molecule] = molecule_count_dict_with_injecting_mismatchable_residue[next_molecule] - 1
			molecule_count = molecule_count - 1
			next_molecule_length = get_molecule_length_from_name(next_molecule)
			right_occupied_length = right_occupied_length + next_molecule_length

			first_layer_right_half_molecule_list.append(next_molecule)

			current_molecule_name = next_molecule

		if right_occupied_length==right_total_length:
			first_layer_right_half_ordered_molecules = first_layer_right_half_molecule_list
			#unused_molecule_count_dict_with_mismatchable_residue = molecule_count_dict_with_injecting_mismatchable_residue
		elif right_occupied_length>right_total_length:
			#[first_layer_ordered_molecules, unused_molecule_count_dict_with_mismatchable_residue] = clean_tail_fill_in_gap(D_site_prob, occupied_length, surface_length, first_layer_molecule_list, molecule_count_dict_with_injecting_mismatchable_residue, molecule_count)
			[first_layer_right_half_ordered_molecules, molecule_count_dict_with_injecting_mismatchable_residue, molecule_count] = clean_tail_fill_in_gap(D_site_prob, right_occupied_length, right_total_length, first_layer_right_half_molecule_list, molecule_count_dict_with_injecting_mismatchable_residue, molecule_count)

		else:
			print("Error: Loop incomplete!!!")
			sys.exit()


	first_layer_ordered_molecules = []
	for kk in range(len(first_layer_left_half_ordered_molecules)-1):
		first_layer_ordered_molecules.append(first_layer_left_half_ordered_molecules[-1-kk])
	#first_layer_ordered_molecules.append(first_molecule)
	for kk in first_layer_right_half_ordered_molecules:
		first_layer_ordered_molecules.append(kk)

	unused_molecule_count_dict_with_mismatchable_residue = molecule_count_dict_with_injecting_mismatchable_residue

	return([first_layer_ordered_molecules, unused_molecule_count_dict_with_mismatchable_residue])


def split_first_layer_molecules_and_assign_to_sites(first_layer_randrev_molecules):
	first_layer_randrev_molecules_split = []
	for molecule in first_layer_randrev_molecules:
		if molecule[0]=='v':#monomer or polymer
			if molecule[4]=='v':#monomer
				first_layer_randrev_molecules_split.append(molecule)
			else:#polymer
				if molecule[1]=='x':#forward
					molecule_comma = molecule.replace('yx','y,x')
				else:#reverse
					molecule_comma = molecule.replace('xy','x,y')
				molecule_split = molecule_comma.split(',')
				for residue in molecule_split:
					first_layer_randrev_molecules_split.append(residue)

		else:#Fz, Gz, E, W
			first_layer_randrev_molecules_split.append(molecule)

	return(first_layer_randrev_molecules_split)


def get_first_layer_attachable_symbol_for_free_monomer(monomer_label):
	attachable_dict = {
	#forward direction
	'vxAyv':[                            'vyBxv','yBxv','vyBx','yBx','vyMxv','yMxv','vyMx','yMx','vyNxv','yNxv','vyNx','yNx'],
	'vxByv':['vyAxv','yAxv','vyAx','yAx',                            'vyMxv','yMxv','vyMx','yMx','vyNxv','yNxv','vyNx','yNx'],
	'vxMyv':['vyAxv','yAxv','vyAx','yAx','vyBxv','yBxv','vyBx','yBx','vyMxv','yMxv','vyMx','yMx','vyNxv','yNxv','vyNx','yNx'],
	'vxNyv':['vyAxv','yAxv','vyAx','yAx','vyBxv','yBxv','vyBx','yBx','vyMxv','yMxv','vyMx','yMx','vyNxv','yNxv','vyNx','yNx'],
	#reverse direction
	'vyAxv':[                            'vxByv','xByv','vxBy','xBy','vxMyv','xMyv','vxMy','xMy','vxNyv','xNyv','vxNy','xNy'],
	'vyBxv':['vxAyv','xAyv','vxAy','xAy',                            'vxMyv','xMyv','vxMy','xMy','vxNyv','xNyv','vxNy','xNy'],
	'vyMxv':['vxAyv','xAyv','vxAy','xAy','vxByv','xByv','vxBy','xBy','vxMyv','xMyv','vxMy','xMy','vxNyv','xNyv','vxNy','xNy'],
	'vyNxv':['vxAyv','xAyv','vxAy','xAy','vxByv','xByv','vxBy','xBy','vxMyv','xMyv','vxMy','xMy','vxNyv','xNyv','vxNy','xNy']
	}
	attachable_first_layer_symbol_list = attachable_dict[monomer_label]
	return(attachable_first_layer_symbol_list)


def get_first_layer_attachable_region_start_index_for_free_polymer(polymer_label, polymer_length, first_layer_randrev_molecules_split):
	v_symbol_stripped_polymer_label = polymer_label.strip('v')
	if v_symbol_stripped_polymer_label[0]=='x':#forward
		molecule_comma = v_symbol_stripped_polymer_label.replace('yx','y,x')
	else:#reverse
		molecule_comma = v_symbol_stripped_polymer_label.replace('xy','x,y')
	v_symbol_stripped_polymer_label_split = molecule_comma.split(',')

	attachable_dict = {
	#forward direction
	'xAy':[                            'vyBxv','yBxv','vyBx','yBx','vyMxv','yMxv','vyMx','yMx','vyNxv','yNxv','vyNx','yNx'],
	'xBy':['vyAxv','yAxv','vyAx','yAx',                            'vyMxv','yMxv','vyMx','yMx','vyNxv','yNxv','vyNx','yNx'],
	'xMy':['vyAxv','yAxv','vyAx','yAx','vyBxv','yBxv','vyBx','yBx','vyMxv','yMxv','vyMx','yMx','vyNxv','yNxv','vyNx','yNx'],
	'xNy':['vyAxv','yAxv','vyAx','yAx','vyBxv','yBxv','vyBx','yBx','vyMxv','yMxv','vyMx','yMx','vyNxv','yNxv','vyNx','yNx'],
	#reverse direction
	'yAx':[                            'vxByv','xByv','vxBy','xBy','vxMyv','xMyv','vxMy','xMy','vxNyv','xNyv','vxNy','xNy'],
	'yBx':['vxAyv','xAyv','vxAy','xAy',                            'vxMyv','xMyv','vxMy','xMy','vxNyv','xNyv','vxNy','xNy'],
	'yMx':['vxAyv','xAyv','vxAy','xAy','vxByv','xByv','vxBy','xBy','vxMyv','xMyv','vxMy','xMy','vxNyv','xNyv','vxNy','xNy'],
	'yNx':['vxAyv','xAyv','vxAy','xAy','vxByv','xByv','vxBy','xBy','vxMyv','xMyv','vxMy','xMy','vxNyv','xNyv','vxNy','xNy']
	}

	attachable_first_layer_region_start_list = []
	for i in range(len(first_layer_randrev_molecules_split)-polymer_length+1):
		region_start_inclusive = i
		region_end_exclusive = i+polymer_length
		first_layer_symbols = first_layer_randrev_molecules_split[region_start_inclusive:region_end_exclusive]
		match_indicator = 'Yes'
		for j in range(polymer_length):
			free_residue = v_symbol_stripped_polymer_label_split[j]
			first_layer_site = first_layer_symbols[j]
			attachable_site_candidates = attachable_dict[free_residue]
			if first_layer_site not in attachable_site_candidates:# must all match
				match_indicator = 'No'
				break
		if match_indicator=='Yes':
			attachable_first_layer_region_start_list.append(region_start_inclusive)

	return(attachable_first_layer_region_start_list)


def get_possible_second_layer_attachment_dict(first_layer_randrev_molecules_split, unused_molecule_count_dict_with_mismatchable_residue):
	possible_second_layer_attachment_dict = {}
	for unused_molecule in unused_molecule_count_dict_with_mismatchable_residue:
		if unused_molecule[0]=='v':#monomer or polymer
			weight = unused_molecule_count_dict_with_mismatchable_residue[unused_molecule]
			if unused_molecule[4]=='v':#monomer
				forward_direction_monomer_label = unused_molecule
				attachable_first_layer_symbol_list = get_first_layer_attachable_symbol_for_free_monomer(forward_direction_monomer_label)
				for i in range(len(first_layer_randrev_molecules_split)):
					first_layer_site = first_layer_randrev_molecules_split[i]
					if first_layer_site in attachable_first_layer_symbol_list:
						attachment_description = unused_molecule + '_forward_attaches_region_' + str(i) + '_to_' +str(i)#inclusive
						product = forward_direction_monomer_label
						possible_second_layer_attachment_dict[attachment_description] = [product, weight]

				reverse_direction_monomer_label = get_reverse_seq(unused_molecule)
				attachable_first_layer_symbol_list = get_first_layer_attachable_symbol_for_free_monomer(reverse_direction_monomer_label)
				for i in range(len(first_layer_randrev_molecules_split)):
					first_layer_site = first_layer_randrev_molecules_split[i]
					if first_layer_site in attachable_first_layer_symbol_list:
						attachment_description = unused_molecule + '_reverse_attaches_region_' + str(i) + '_to_' +str(i)#inclusive
						product = reverse_direction_monomer_label
						possible_second_layer_attachment_dict[attachment_description] = [product, weight]

			else:#polymer
				polymer_length = get_molecule_length_from_name(unused_molecule)
				#forward direction
				forward_direction_polymer_label = unused_molecule
				attachable_first_layer_region_start_list = get_first_layer_attachable_region_start_index_for_free_polymer(forward_direction_polymer_label, polymer_length, first_layer_randrev_molecules_split)
				for attachable_first_layer_region_start in attachable_first_layer_region_start_list:
					attachment_description = unused_molecule + '_forward_attaches_region_' + str(attachable_first_layer_region_start) + '_to_' +str(attachable_first_layer_region_start+polymer_length-1)#inclusive
					product = forward_direction_polymer_label
					possible_second_layer_attachment_dict[attachment_description] = [product, weight]

				reverse_direction_polymer_label = get_reverse_seq(unused_molecule)
				attachable_first_layer_region_start_list = get_first_layer_attachable_region_start_index_for_free_polymer(reverse_direction_polymer_label, polymer_length, first_layer_randrev_molecules_split)
				for attachable_first_layer_region_start in attachable_first_layer_region_start_list:
					attachment_description = unused_molecule + '_reverse_attaches_region_' + str(attachable_first_layer_region_start) + '_to_' +str(attachable_first_layer_region_start+polymer_length-1)#inclusive
					product = reverse_direction_polymer_label
					possible_second_layer_attachment_dict[attachment_description] = [product, weight]

	return(possible_second_layer_attachment_dict)


def check_if_candidate_region_overlaps_picked_region(affected_region_start_inclusive, affected_region_end_inclusive, candidate_affected_region_start_inclusive, candidate_affected_region_end_inclusive):
	if affected_region_end_inclusive<candidate_affected_region_start_inclusive:
		candidate_region_overlaps_picked_region = 'No'
	elif affected_region_start_inclusive>candidate_affected_region_end_inclusive:
		candidate_region_overlaps_picked_region = 'No'
	else:
		candidate_region_overlaps_picked_region = 'Yes'
	return(candidate_region_overlaps_picked_region)


def update_surface_profile_and_attachment_dict_and_unused_molecule_dict(picked_attachment, fully_adsorption_surface_profile, possible_second_layer_attachment_dict, unused_molecule_count_dict_with_mismatchable_residue):
	attach_molecule_label = picked_attachment.split('_')[0]
	affected_region_description = picked_attachment.split('_region_')[1]
	affected_region_start_inclusive = int(affected_region_description.split('_to_')[0])
	affected_region_end_inclusive = int(affected_region_description.split('_to_')[1])
	attachment_product = possible_second_layer_attachment_dict[picked_attachment][0]
	if affected_region_start_inclusive==affected_region_end_inclusive:# this is a monomer
		fully_adsorption_surface_profile[affected_region_start_inclusive].append(attachment_product)
	else:# this is a polymer
		if attachment_product[1]=='x':#forward
			product_comma = attachment_product.replace('yx','y,x')
		else:#reverse
			product_comma = attachment_product.replace('xy','x,y')
		product_split = product_comma.split(',')
		product_length = len(product_split)
		for m in range(product_length):
			fully_adsorption_surface_profile[affected_region_start_inclusive+m].append(product_split[m])

	unused_molecule_count_dict_with_mismatchable_residue[attach_molecule_label] = unused_molecule_count_dict_with_mismatchable_residue[attach_molecule_label] - 1

	for attachment_description in possible_second_layer_attachment_dict:
		candidate_affected_region_description = attachment_description.split('_region_')[1]
		candidate_affected_region_start_inclusive = int(candidate_affected_region_description.split('_to_')[0])
		candidate_affected_region_end_inclusive = int(candidate_affected_region_description.split('_to_')[1])
		candidate_region_overlaps_picked_region = check_if_candidate_region_overlaps_picked_region(affected_region_start_inclusive, affected_region_end_inclusive, candidate_affected_region_start_inclusive, candidate_affected_region_end_inclusive)
		if candidate_region_overlaps_picked_region=='Yes':
			possible_second_layer_attachment_dict[attachment_description][1] = 0# if the regions overlap, this attachment is no longer available

		candidate_molecule_label = attachment_description.split('_')[0]
		if candidate_molecule_label==attach_molecule_label:
			if unused_molecule_count_dict_with_mismatchable_residue[candidate_molecule_label]==0:
				possible_second_layer_attachment_dict[attachment_description][1] = 0# if no molecule is available, this attachment is impossible

	return([fully_adsorption_surface_profile, possible_second_layer_attachment_dict, unused_molecule_count_dict_with_mismatchable_residue])


def attach_second_layer(surface_length, first_layer_randrev_molecules_split, possible_second_layer_attachment_dict, unused_molecule_count_dict_with_mismatchable_residue):
	fully_adsorption_surface_profile = []
	for k in range(surface_length):
		fully_adsorption_surface_profile.append([first_layer_randrev_molecules_split[k]])

	sum_weight = 0
	attachment_description_list = []
	for attachment_description in possible_second_layer_attachment_dict:
		weight = possible_second_layer_attachment_dict[attachment_description][1]
		sum_weight = sum_weight + weight
		attachment_description_list.append(attachment_description)

	while sum_weight>0:
		p_list = []
		for attachment_description in attachment_description_list:
			prob = possible_second_layer_attachment_dict[attachment_description][1]/sum_weight
			p_list.append(prob)
		picked_attachment = np.random.choice(attachment_description_list, p=p_list)
		[fully_adsorption_surface_profile, possible_second_layer_attachment_dict, unused_molecule_count_dict_with_mismatchable_residue] = update_surface_profile_and_attachment_dict_and_unused_molecule_dict(picked_attachment, fully_adsorption_surface_profile, possible_second_layer_attachment_dict, unused_molecule_count_dict_with_mismatchable_residue)
		sum_weight = 0
		for attachment_description in attachment_description_list:
			weight = possible_second_layer_attachment_dict[attachment_description][1]
			sum_weight = sum_weight + weight

	return([fully_adsorption_surface_profile, unused_molecule_count_dict_with_mismatchable_residue])


def Stochastic_cover_surface_with_molecules_until_saturation(molecule_name_list, surface_length, mismatch_prob, D_site_prob):# concentration driven only
	molecule_count = len(molecule_name_list)

	molecule_name_with_polarity_list = []
	for mol in molecule_name_list:
		if 'A' in mol:
			mol_polar = convert_normal_polymer_name_to_xy_name(mol)
			molecule_name_with_polarity_list.append(mol_polar)
		elif 'B' in mol:
			mol_polar = convert_normal_polymer_name_to_xy_name(mol)
			molecule_name_with_polarity_list.append(mol_polar)
		elif mol=='F':
			molecule_name_with_polarity_list.append('Fz')
		elif mol=='G':
			molecule_name_with_polarity_list.append('Gz')
		else:
			molecule_name_with_polarity_list.append(mol)

	molecule_name_list_after_injecting_mismatchable_residue = inject_mismatch_residue(molecule_name_with_polarity_list, mismatch_prob)
	molecule_count_dict_with_injecting_mismatchable_residue = convert_list_to_count_dict(molecule_name_list_after_injecting_mismatchable_residue)

	[first_layer_ordered_molecules, unused_molecule_count_dict_with_mismatchable_residue] = fill_in_first_layer_with_molecules(surface_length, D_site_prob, molecule_count_dict_with_injecting_mismatchable_residue, molecule_count)
	first_layer_randrev_molecules = []
	for mol in first_layer_ordered_molecules:
		randrev_mol = randomly_reverse_molecule(mol)
		first_layer_randrev_molecules.append(randrev_mol)

	#unused_E_count = unused_molecule_count_dict_with_mismatchable_residue['E']
	#unused_W_count = unused_molecule_count_dict_with_mismatchable_residue['W']
	#unused_Fz_count = unused_molecule_count_dict_with_mismatchable_residue['Fz']
	#unused_Gz_count = unused_molecule_count_dict_with_mismatchable_residue['Gz']

	first_layer_randrev_molecules_split = split_first_layer_molecules_and_assign_to_sites(first_layer_randrev_molecules)

#=======================================
#	print(*first_layer_randrev_molecules_split, sep='\t')
#===========================================

	possible_second_layer_attachment_dict = get_possible_second_layer_attachment_dict(first_layer_randrev_molecules_split, unused_molecule_count_dict_with_mismatchable_residue)

	[fully_adsorption_surface_profile, unused_molecule_count_dict_with_mismatchable_residue] = attach_second_layer(surface_length, first_layer_randrev_molecules_split, possible_second_layer_attachment_dict, unused_molecule_count_dict_with_mismatchable_residue)

	return([fully_adsorption_surface_profile, unused_molecule_count_dict_with_mismatchable_residue])


def remove_MN_labels_from_surface_profile(fully_adsorption_surface_profile):
	for i in range(len(fully_adsorption_surface_profile)):
		site = fully_adsorption_surface_profile[i]
		for j in range(len(site)):
			symbol = site[j]
			symbol = symbol.replace('M','A')
			symbol = symbol.replace('N','B')
			fully_adsorption_surface_profile[i][j] = symbol
	return(fully_adsorption_surface_profile)


def remove_MN_labels_from_free_molecules(unused_molecule_count_dict_with_mismatchable_residue):
	unused_molecule_count_dict_normal_name = {}
	for xy_name in unused_molecule_count_dict_with_mismatchable_residue:
		if unused_molecule_count_dict_with_mismatchable_residue[xy_name]>0:
			if 'v' in xy_name:#monomers and polymers
				#xy_name = orient_polymer_seq(xy_name)
				normal_name = convert_xy_name_to_normal_polymer_name(xy_name)
				normal_name = normal_name.replace('M','A')
				normal_name = normal_name.replace('N','B')
				if normal_name not in unused_molecule_count_dict_normal_name:
					unused_molecule_count_dict_normal_name[normal_name] = unused_molecule_count_dict_with_mismatchable_residue[xy_name]
				else:
					unused_molecule_count_dict_normal_name[normal_name] = unused_molecule_count_dict_normal_name[normal_name] + unused_molecule_count_dict_with_mismatchable_residue[xy_name]
			
			elif 'z' in xy_name:#Fz,Gz
				normal_name = xy_name.replace('z','')
				unused_molecule_count_dict_normal_name[normal_name] = unused_molecule_count_dict_with_mismatchable_residue[xy_name]

			else:#E,W
				unused_molecule_count_dict_normal_name[xy_name] = unused_molecule_count_dict_with_mismatchable_residue[xy_name]

	return(unused_molecule_count_dict_normal_name)


def scan_borders_and_get_reaction_types(fully_adsorption_surface_profile):
	kination_deactivation_border_list = []
	ligation_breakage_border_list = []
	for border_id in range(len(fully_adsorption_surface_profile)-1):

		left_site = fully_adsorption_surface_profile[border_id]
		right_site = fully_adsorption_surface_profile[border_id+1]
		#left_layer_num=len(left_site)
		#right_layer_num = len(right_site)
#=============================================
		#zip_left_right = [left_site, right_site]

# Fz + E ==> vxAyv + W
# If a reaction occurs between W and vyA/Bxv on the other side, this site will become Fz~E

# vxAyv + W ==> Fz + E
# If a reaction occurs between E and zF/G on the other side, this site will become vxAyv~W

# E + zF ==> W + vyAxv
# If a reaction occurs between vxA/Byv and W on the other side, this site will become E~zF

# W + vyAxv ==> E + zF
# If a reaction occurs between F/Gz and E on the other side, this site will become W~vyAxv

# Gz + E ==> vxByv + W
# vxByv + W ==> Gz + E
# E + zG ==> W + vyBxv
# W + vyBxv ==> E + zG

# Note that indirect reaction sites are considered when computing possible borders, but whether a reaction can occur will be checked whenever a relevant collision is stochastically picked.
		if left_site in [['E'],['W']]:
			if right_site in [['zF'],['zG'],['vyAxv'],['vyBxv']]:
				kination_deactivation_border_list.append(border_id)
		elif left_site in [['Fz'],['Gz'],['vxAyv'],['vxByv']]:
			if right_site in [['E'],['W']]:
				kination_deactivation_border_list.append(border_id)

#=============================================

		elif len(left_site)+len(right_site)==4:
			left_site_top_layer_right_edge = left_site[1][-2:]
			right_site_top_layer_left_edge = right_site[1][0:2]
			zip_left_right_2nd_layer_TwoLetters = [left_site_top_layer_right_edge, right_site_top_layer_left_edge]
			zip_left_right_2nd_layer_OneLetter = [left_site_top_layer_right_edge[1], right_site_top_layer_left_edge[0]]
# ~yv + vx~ ==> ~yx~
			if zip_left_right_2nd_layer_TwoLetters == ['yv','vx']:
				ligation_breakage_border_list.append(border_id)
# ~yx~ ==> ~yv + vx~
			elif zip_left_right_2nd_layer_OneLetter == ['y','x']:
				ligation_breakage_border_list.append(border_id)
# ~xv + vy~ ==> ~xy~
			elif zip_left_right_2nd_layer_TwoLetters == ['xv','vy']:
				ligation_breakage_border_list.append(border_id)
# ~xy~ ==> ~xv + vy~
			elif zip_left_right_2nd_layer_OneLetter == ['x','y']:
				ligation_breakage_border_list.append(border_id)

	return([kination_deactivation_border_list, ligation_breakage_border_list])


def calculate_numbers_of_collisions(base_collision_number_per_residue_per_cycle, unused_molecule_count_dict_normal_name, kinase_dict, ligase_dict, volume, natural_kination_catalyst_collisions_per_volume_per_cycle, natural_ligation_catalyst_collisions_per_volume_per_cycle):
	# Get the types and counts of catalysts
	kinase_collision_number_dict = {}
	ligase_collision_number_dict = {}
	for normal_name in unused_molecule_count_dict_normal_name:
		if normal_name in kinase_dict:
			#present_kinase_label_list.append(normal_name)
			#kinase_collision_number_dict[normal_name] = int(base_collision_number_per_residue_per_cycle * unused_molecule_count_dict_normal_name[normal_name] / volume)
			enzyme_size = len(normal_name)
			kinase_collision_number_dict[normal_name] = base_collision_number_per_residue_per_cycle * unused_molecule_count_dict_normal_name[normal_name] / (enzyme_size**0.5)
		elif normal_name in ligase_dict:#one polymer must either be a kinase or a ligase !!!!!!!
			#present_ligase_label_list.append(normal_name)
			#ligase_collision_number_dict[normal_name] = int(base_collision_number_per_residue_per_cycle * unused_molecule_count_dict_normal_name[normal_name] / volume)
			enzyme_size = len(normal_name)
			ligase_collision_number_dict[normal_name] = base_collision_number_per_residue_per_cycle * unused_molecule_count_dict_normal_name[normal_name] / (enzyme_size**0.5)

	total_kinase_collision_number = 0
	for i in kinase_collision_number_dict:
		total_kinase_collision_number = total_kinase_collision_number + kinase_collision_number_dict[i]
	
	total_ligase_collision_number = 0
	for j in ligase_collision_number_dict:
		total_ligase_collision_number = total_ligase_collision_number + ligase_collision_number_dict[j]

	total_natural_kination_catalyst_collision_number = natural_kination_catalyst_collisions_per_volume_per_cycle * volume
	total_natural_ligation_catalyst_collision_number = natural_ligation_catalyst_collisions_per_volume_per_cycle * volume

	return([kinase_collision_number_dict, ligase_collision_number_dict, total_kinase_collision_number, total_ligase_collision_number, total_natural_kination_catalyst_collision_number, total_natural_ligation_catalyst_collision_number])


def get_result_after_kinase_finds_target(border_id, kinase_seq, fully_adsorption_surface_profile, kinase_dict, kination_RF_ratio):
	left_site = fully_adsorption_surface_profile[border_id]
	right_site = fully_adsorption_surface_profile[border_id+1]

	successful_kination_prob = kinase_dict[kinase_seq]
	successful_deactivation_prob = successful_kination_prob*kination_RF_ratio

	reaction_type = 'No_reaction'
	zip_left_right = [left_site, right_site]
#=============================================
# Here, indirect sites (e.g., Fz~W, vxAyv~E) are not considered. 
# Fz + E ==> vxAyv + W
	if zip_left_right == [['Fz'],['E']]:
		reaction_type = 'kination'
		reaction_proceeds = np.random.choice(['Yes','No'], p=[successful_kination_prob, 1-successful_kination_prob])
		if reaction_proceeds=='Yes':
			fully_adsorption_surface_profile[border_id] = ['vxAyv']
			fully_adsorption_surface_profile[border_id+1] = ['W']

# vxAyv + W ==> Fz + E
	elif zip_left_right == [['vxAyv'],['W']]:
		reaction_type = 'deactivation'
		reaction_proceeds = np.random.choice(['Yes','No'], p=[successful_deactivation_prob, 1-successful_deactivation_prob])
		if reaction_proceeds=='Yes':
			fully_adsorption_surface_profile[border_id] = ['Fz']
			fully_adsorption_surface_profile[border_id+1] = ['E']

# E + zF ==> W + vyAxv
	elif zip_left_right == [['E'],['zF']]:
		reaction_type = 'kination'
		reaction_proceeds = np.random.choice(['Yes','No'], p=[successful_kination_prob, 1-successful_kination_prob])
		if reaction_proceeds=='Yes':
			fully_adsorption_surface_profile[border_id] = ['W']
			fully_adsorption_surface_profile[border_id+1] = ['vyAxv']

# W + vyAxv ==> E + zF
	elif zip_left_right == [['W'],['vyAxv']]:
		reaction_type = 'deactivation'
		reaction_proceeds = np.random.choice(['Yes','No'], p=[successful_deactivation_prob, 1-successful_deactivation_prob])
		if reaction_proceeds=='Yes':
			fully_adsorption_surface_profile[border_id] = ['E']
			fully_adsorption_surface_profile[border_id+1] = ['zF']

#=============================================
# Gz + E ==> vxByv + W
	elif zip_left_right == [['Gz'],['E']]:
		reaction_type = 'kination'
		reaction_proceeds = np.random.choice(['Yes','No'], p=[successful_kination_prob, 1-successful_kination_prob])
		if reaction_proceeds=='Yes':
			fully_adsorption_surface_profile[border_id] = ['vxByv']
			fully_adsorption_surface_profile[border_id+1] = ['W']

# vxByv + W ==> Gz + E
	elif zip_left_right == [['vxByv'],['W']]:
		reaction_type = 'deactivation'
		reaction_proceeds = np.random.choice(['Yes','No'], p=[successful_deactivation_prob, 1-successful_deactivation_prob])
		if reaction_proceeds=='Yes':
			fully_adsorption_surface_profile[border_id] = ['Gz']
			fully_adsorption_surface_profile[border_id+1] = ['E']

# E + zG ==> W + vyBxv
	elif zip_left_right == [['E'],['zG']]:
		reaction_type = 'kination'
		reaction_proceeds = np.random.choice(['Yes','No'], p=[successful_kination_prob, 1-successful_kination_prob])
		if reaction_proceeds=='Yes':
			fully_adsorption_surface_profile[border_id] = ['W']
			fully_adsorption_surface_profile[border_id+1] = ['vyBxv']

# W + vyBxv ==> E + zG
	elif zip_left_right == [['W'],['vyBxv']]:
		reaction_type = 'deactivation'
		reaction_proceeds = np.random.choice(['Yes','No'], p=[successful_deactivation_prob, 1-successful_deactivation_prob])
		if reaction_proceeds=='Yes':
			fully_adsorption_surface_profile[border_id] = ['E']
			fully_adsorption_surface_profile[border_id+1] = ['zG']

	return([reaction_type, fully_adsorption_surface_profile])


def get_result_after_ligase_finds_target(border_id, ligase_seq, fully_adsorption_surface_profile, ligase_dict, ligation_RF_ratio):
	left_site = fully_adsorption_surface_profile[border_id]
	right_site = fully_adsorption_surface_profile[border_id+1]

	left_site_top_layer_right_edge = left_site[1][-2:]
	right_site_top_layer_left_edge = right_site[1][0:2]

	successful_ligation_prob = ligase_dict[ligase_seq]
	successful_breakage_prob = successful_ligation_prob*ligation_RF_ratio

	reaction_type = 'No_reaction'
	zip_left_right_2nd_layer_TwoLetters = [left_site_top_layer_right_edge, right_site_top_layer_left_edge]
	zip_left_right_2nd_layer_OneLetter = [left_site_top_layer_right_edge[1], right_site_top_layer_left_edge[0]]
#=============================================
# ~yv + vx~ ==> ~yx~
	if zip_left_right_2nd_layer_TwoLetters == ['yv','vx']:
		reaction_type = 'ligation'
		reaction_proceeds = np.random.choice(['Yes','No'], p=[successful_ligation_prob, 1-successful_ligation_prob])
		if reaction_proceeds=='Yes':
			fully_adsorption_surface_profile[border_id][1] = fully_adsorption_surface_profile[border_id][1][0:-1]
			fully_adsorption_surface_profile[border_id+1][1] = fully_adsorption_surface_profile[border_id+1][1][1:]

# ~yx~ ==> ~yv + vx~
	elif zip_left_right_2nd_layer_OneLetter == ['y','x']:
		reaction_type = 'breakage'
		reaction_proceeds = np.random.choice(['Yes','No'], p=[successful_breakage_prob, 1-successful_breakage_prob])
		if reaction_proceeds=='Yes':
			fully_adsorption_surface_profile[border_id][1] = fully_adsorption_surface_profile[border_id][1]+'v'
			fully_adsorption_surface_profile[border_id+1][1] = 'v'+fully_adsorption_surface_profile[border_id+1][1]

# ~xv + vy~ ==> ~xy~
	elif zip_left_right_2nd_layer_TwoLetters == ['xv','vy']:
		reaction_type = 'ligation'
		reaction_proceeds = np.random.choice(['Yes','No'], p=[successful_ligation_prob, 1-successful_ligation_prob])
		if reaction_proceeds=='Yes':
			fully_adsorption_surface_profile[border_id][1] = fully_adsorption_surface_profile[border_id][1][0:-1]
			fully_adsorption_surface_profile[border_id+1][1] = fully_adsorption_surface_profile[border_id+1][1][1:]

# ~xy~ ==> ~xv + vy~
	elif zip_left_right_2nd_layer_OneLetter == ['x','y']:
		reaction_type = 'breakage'
		reaction_proceeds = np.random.choice(['Yes','No'], p=[successful_breakage_prob, 1-successful_breakage_prob])
		if reaction_proceeds=='Yes':
			fully_adsorption_surface_profile[border_id][1] = fully_adsorption_surface_profile[border_id][1]+'v'
			fully_adsorption_surface_profile[border_id+1][1] = 'v'+fully_adsorption_surface_profile[border_id+1][1]

	return([reaction_type, fully_adsorption_surface_profile])


def get_result_after_natural_kination_catalyst_finds_target(border_id, fully_adsorption_surface_profile, spontaneous_kination_prob, kination_RF_ratio):
	left_site = fully_adsorption_surface_profile[border_id]
	right_site = fully_adsorption_surface_profile[border_id+1]

	spontaneous_deactivation_prob = spontaneous_kination_prob*kination_RF_ratio

	reaction_type = 'No_reaction'
	zip_left_right = [left_site, right_site]
#=============================================
# Here, indirect sites (e.g., Fz~W, vxAyv~E) are not considered. 
# Fz + E ==> vxAyv + W
	if zip_left_right == [['Fz'],['E']]:
		reaction_type = 'kination'
		reaction_proceeds = np.random.choice(['Yes','No'], p=[spontaneous_kination_prob, 1-spontaneous_kination_prob])
		if reaction_proceeds=='Yes':
			fully_adsorption_surface_profile[border_id] = ['vxAyv']
			fully_adsorption_surface_profile[border_id+1] = ['W']

# vxAyv + W ==> Fz + E
	elif zip_left_right == [['vxAyv'],['W']]:
		reaction_type = 'deactivation'
		reaction_proceeds = np.random.choice(['Yes','No'], p=[spontaneous_deactivation_prob, 1-spontaneous_deactivation_prob])
		if reaction_proceeds=='Yes':
			fully_adsorption_surface_profile[border_id] = ['Fz']
			fully_adsorption_surface_profile[border_id+1] = ['E']

# E + zF ==> W + vyAxv
	elif zip_left_right == [['E'],['zF']]:
		reaction_type = 'kination'
		reaction_proceeds = np.random.choice(['Yes','No'], p=[spontaneous_kination_prob, 1-spontaneous_kination_prob])
		if reaction_proceeds=='Yes':
			fully_adsorption_surface_profile[border_id] = ['W']
			fully_adsorption_surface_profile[border_id+1] = ['vyAxv']

# W + vyAxv ==> E + zF
	elif zip_left_right == [['W'],['vyAxv']]:
		reaction_type = 'deactivation'
		reaction_proceeds = np.random.choice(['Yes','No'], p=[spontaneous_deactivation_prob, 1-spontaneous_deactivation_prob])
		if reaction_proceeds=='Yes':
			fully_adsorption_surface_profile[border_id] = ['E']
			fully_adsorption_surface_profile[border_id+1] = ['zF']

#=============================================
# Gz + E ==> vxByv + W
	elif zip_left_right == [['Gz'],['E']]:
		reaction_type = 'kination'
		reaction_proceeds = np.random.choice(['Yes','No'], p=[spontaneous_kination_prob, 1-spontaneous_kination_prob])
		if reaction_proceeds=='Yes':
			fully_adsorption_surface_profile[border_id] = ['vxByv']
			fully_adsorption_surface_profile[border_id+1] = ['W']

# vxByv + W ==> Gz + E
	elif zip_left_right == [['vxByv'],['W']]:
		reaction_type = 'deactivation'
		reaction_proceeds = np.random.choice(['Yes','No'], p=[spontaneous_deactivation_prob, 1-spontaneous_deactivation_prob])
		if reaction_proceeds=='Yes':
			fully_adsorption_surface_profile[border_id] = ['Gz']
			fully_adsorption_surface_profile[border_id+1] = ['E']

# E + zG ==> W + vyBxv
	elif zip_left_right == [['E'],['zG']]:
		reaction_type = 'kination'
		reaction_proceeds = np.random.choice(['Yes','No'], p=[spontaneous_kination_prob, 1-spontaneous_kination_prob])
		if reaction_proceeds=='Yes':
			fully_adsorption_surface_profile[border_id] = ['W']
			fully_adsorption_surface_profile[border_id+1] = ['vyBxv']

# W + vyBxv ==> E + zG
	elif zip_left_right == [['W'],['vyBxv']]:
		reaction_type = 'deactivation'
		reaction_proceeds = np.random.choice(['Yes','No'], p=[spontaneous_deactivation_prob, 1-spontaneous_deactivation_prob])
		if reaction_proceeds=='Yes':
			fully_adsorption_surface_profile[border_id] = ['E']
			fully_adsorption_surface_profile[border_id+1] = ['zG']

	return([reaction_type, fully_adsorption_surface_profile])


def get_result_after_natural_ligation_catalyst_finds_target(border_id, fully_adsorption_surface_profile, spontaneous_ligation_prob, ligation_RF_ratio):
	left_site = fully_adsorption_surface_profile[border_id]
	right_site = fully_adsorption_surface_profile[border_id+1]

	left_site_top_layer_right_edge = left_site[1][-2:]
	right_site_top_layer_left_edge = right_site[1][0:2]

	spontaneous_breakage_prob = spontaneous_ligation_prob*ligation_RF_ratio

	reaction_type = 'No_reaction'
	zip_left_right_2nd_layer_TwoLetters = [left_site_top_layer_right_edge, right_site_top_layer_left_edge]
	zip_left_right_2nd_layer_OneLetter = [left_site_top_layer_right_edge[1], right_site_top_layer_left_edge[0]]
#=============================================
# ~yv + vx~ ==> ~yx~
	if zip_left_right_2nd_layer_TwoLetters == ['yv','vx']:
		reaction_type = 'ligation'
		reaction_proceeds = np.random.choice(['Yes','No'], p=[spontaneous_ligation_prob, 1-spontaneous_ligation_prob])
		if reaction_proceeds=='Yes':
			fully_adsorption_surface_profile[border_id][1] = fully_adsorption_surface_profile[border_id][1][0:-1]
			fully_adsorption_surface_profile[border_id+1][1] = fully_adsorption_surface_profile[border_id+1][1][1:]

# ~yx~ ==> ~yv + vx~
	elif zip_left_right_2nd_layer_OneLetter == ['y','x']:
		reaction_type = 'breakage'
		reaction_proceeds = np.random.choice(['Yes','No'], p=[spontaneous_breakage_prob, 1-spontaneous_breakage_prob])
		if reaction_proceeds=='Yes':
			fully_adsorption_surface_profile[border_id][1] = fully_adsorption_surface_profile[border_id][1]+'v'
			fully_adsorption_surface_profile[border_id+1][1] = 'v'+fully_adsorption_surface_profile[border_id+1][1]

# ~xv + vy~ ==> ~xy~
	elif zip_left_right_2nd_layer_TwoLetters == ['xv','vy']:
		reaction_type = 'ligation'
		reaction_proceeds = np.random.choice(['Yes','No'], p=[spontaneous_ligation_prob, 1-spontaneous_ligation_prob])
		if reaction_proceeds=='Yes':
			fully_adsorption_surface_profile[border_id][1] = fully_adsorption_surface_profile[border_id][1][0:-1]
			fully_adsorption_surface_profile[border_id+1][1] = fully_adsorption_surface_profile[border_id+1][1][1:]

# ~xy~ ==> ~xv + vy~
	elif zip_left_right_2nd_layer_OneLetter == ['x','y']:
		reaction_type = 'breakage'
		reaction_proceeds = np.random.choice(['Yes','No'], p=[spontaneous_breakage_prob, 1-spontaneous_breakage_prob])
		if reaction_proceeds=='Yes':
			fully_adsorption_surface_profile[border_id][1] = fully_adsorption_surface_profile[border_id][1]+'v'
			fully_adsorption_surface_profile[border_id+1][1] = 'v'+fully_adsorption_surface_profile[border_id+1][1]

	return([reaction_type, fully_adsorption_surface_profile])


def semi_Gillespie_determine_final_status(surface_length, fully_adsorption_surface_profile, kination_deactivation_border_list, ligation_breakage_border_list, kinase_collision_number_dict, ligase_collision_number_dict, total_kinase_collision_number, total_ligase_collision_number, kinase_dict, ligase_dict, spontaneous_kination_prob, kination_RF_ratio, spontaneous_ligation_prob, ligation_RF_ratio, total_natural_kination_catalyst_collision_number, total_natural_ligation_catalyst_collision_number):
	total_catalyst_collision_number = total_kinase_collision_number + total_ligase_collision_number + total_natural_kination_catalyst_collision_number + total_natural_ligation_catalyst_collision_number
	if total_catalyst_collision_number<=1e9:
		simulated_catalyst_collision_number = np.random.poisson(lam=int(total_catalyst_collision_number))
	else:
		simulated_catalyst_collision_number = int(np.random.normal(loc=total_catalyst_collision_number, scale=total_catalyst_collision_number**0.5))
		if simulated_catalyst_collision_number<0:
			simulated_catalyst_collision_number = 0

	p_kinase_collision_with_any_borders = total_kinase_collision_number/total_catalyst_collision_number
	p_ligase_collision_with_any_borders = total_ligase_collision_number/total_catalyst_collision_number
	p_natural_kination_catalyst_collision_with_any_borders = total_natural_kination_catalyst_collision_number/total_catalyst_collision_number
	p_natural_ligation_catalyst_collision_with_any_borders = total_natural_ligation_catalyst_collision_number/total_catalyst_collision_number


	p_collision_with_kination_deactivation_borders = len(kination_deactivation_border_list)/(surface_length-1)
	p_collision_with_ligation_breakage_borders = len(ligation_breakage_border_list)/(surface_length-1)

	p_kinase_collision_with_kination_deactivation_borders = p_kinase_collision_with_any_borders * p_collision_with_kination_deactivation_borders
	p_ligase_collision_with_ligation_breakage_borders = p_ligase_collision_with_any_borders * p_collision_with_ligation_breakage_borders
	p_natural_kination_catalyst_collision_with_kination_deactivation_borders = p_natural_kination_catalyst_collision_with_any_borders * p_collision_with_kination_deactivation_borders
	p_natural_ligation_catalyst_collision_with_ligation_breakage_borders = p_natural_ligation_catalyst_collision_with_any_borders * p_collision_with_ligation_breakage_borders

	p_other_collissions = 1 - p_kinase_collision_with_kination_deactivation_borders - p_ligase_collision_with_ligation_breakage_borders - p_natural_kination_catalyst_collision_with_kination_deactivation_borders - p_natural_ligation_catalyst_collision_with_ligation_breakage_borders

	kinase_seq_list = []
	kinase_seq_pick_prob_list = []
	for kinase_seq in kinase_collision_number_dict:
		kinase_seq_list.append(kinase_seq)
		kinase_seq_pick_prob = kinase_collision_number_dict[kinase_seq]/total_kinase_collision_number
		kinase_seq_pick_prob_list.append(kinase_seq_pick_prob)
		#===================Make sure that the sum is 1.
		kinase_seq_pick_prob_list[0] = 1-sum(kinase_seq_pick_prob_list[1:])
		if kinase_seq_pick_prob_list[0]<0:
			kinase_seq_pick_prob_list[0] = 0

	ligase_seq_list = []
	ligase_seq_pick_prob_list = []
	for ligase_seq in ligase_collision_number_dict:
		ligase_seq_list.append(ligase_seq)
		ligase_seq_pick_prob = ligase_collision_number_dict[ligase_seq]/total_ligase_collision_number
		ligase_seq_pick_prob_list.append(ligase_seq_pick_prob)
		#===================Make sure that the sum is 1.
		ligase_seq_pick_prob_list[0] = 1-sum(ligase_seq_pick_prob_list[1:])
		if ligase_seq_pick_prob_list[0]<0:
			ligase_seq_pick_prob_list[0] = 0

	for collision_event in range(simulated_catalyst_collision_number):
		collision_type = np.random.choice(['kinase_finds_target','ligase_finds_target','natural_kination_catalyst_finds_target','natural_ligation_catalyst_finds_target','useless_collision'], p=[p_kinase_collision_with_kination_deactivation_borders, p_ligase_collision_with_ligation_breakage_borders, p_natural_kination_catalyst_collision_with_kination_deactivation_borders, p_natural_ligation_catalyst_collision_with_ligation_breakage_borders, p_other_collissions])
		if collision_type == 'useless_collision':
			reaction_type = 'No_reaction'

		elif collision_type == 'kinase_finds_target':
			border_id = np.random.choice(kination_deactivation_border_list)
			kinase_seq = np.random.choice(kinase_seq_list, p=kinase_seq_pick_prob_list)
			[reaction_type, fully_adsorption_surface_profile] = get_result_after_kinase_finds_target(border_id, kinase_seq, fully_adsorption_surface_profile, kinase_dict, kination_RF_ratio)

		elif collision_type == 'ligase_finds_target':
			border_id = np.random.choice(ligation_breakage_border_list)
			ligase_seq = np.random.choice(ligase_seq_list, p=ligase_seq_pick_prob_list)
			[reaction_type, fully_adsorption_surface_profile] = get_result_after_ligase_finds_target(border_id, ligase_seq, fully_adsorption_surface_profile, ligase_dict, ligation_RF_ratio)

		elif collision_type == 'natural_kination_catalyst_finds_target':
			border_id = np.random.choice(kination_deactivation_border_list)
			[reaction_type, fully_adsorption_surface_profile] = get_result_after_natural_kination_catalyst_finds_target(border_id, fully_adsorption_surface_profile, spontaneous_kination_prob, kination_RF_ratio)

		elif collision_type == 'natural_ligation_catalyst_finds_target':
			border_id = np.random.choice(ligation_breakage_border_list)
			[reaction_type, fully_adsorption_surface_profile] = get_result_after_natural_ligation_catalyst_finds_target(border_id, fully_adsorption_surface_profile, spontaneous_ligation_prob, ligation_RF_ratio)

	return(fully_adsorption_surface_profile)


def release_molecules_from_surface(fully_adsorption_surface_profile, unused_molecule_count_dict_normal_name):
	free_molecules_list = []
	#layer1
	layer1_list = []
	for site in fully_adsorption_surface_profile:
		layer1 = site[0]
		layer1 = layer1.replace('z','')#impossible for layer2
		layer1_list.append(layer1)
	layer1_str = '.'.join(layer1_list)
	#print(layer1_str)
	layer1_str = layer1_str.replace('x.y','xy')
	#print(layer1_str)
	layer1_str = layer1_str.replace('y.x','yx')
	#print(layer1_str)
	if layer1_str!='':
		layer1_mol_list = layer1_str.split('.')
	else:
		layer1_mol_list = []
	#print(layer1_mol_list)
	for mol in layer1_mol_list:
		if 'x' in mol:
			mol = orient_polymer_seq(mol)
			mol = convert_xy_name_to_normal_polymer_name(mol)
		free_molecules_list.append(mol)

	#layer2
	layer2_list = []
	for site in fully_adsorption_surface_profile:
		if len(site)==2:
			layer2 = site[1]
			layer2_list.append(layer2)
	layer2_str = '.'.join(layer2_list)
	#print(layer2_str)
	layer2_str = layer2_str.replace('x.y','xy')
	#print(layer2_str)
	layer2_str = layer2_str.replace('y.x','yx')
	#print(layer2_str)
	if layer2_str!='':
		layer2_mol_list = layer2_str.split('.')
	else:
		layer2_mol_list = []
	#print(layer2_mol_list)
	for mol in layer2_mol_list:
		if 'x' in mol:
			mol = orient_polymer_seq(mol)
			mol = convert_xy_name_to_normal_polymer_name(mol)
		free_molecules_list.append(mol)
	#print(free_molecules_list)

	for mol in unused_molecule_count_dict_normal_name:
		mol_count = unused_molecule_count_dict_normal_name[mol]
		for i in range(mol_count):
			free_molecules_list.append(mol)

	return(free_molecules_list)


def formatted_print_two_layer_profile(fully_adsorption_surface_profile):
	surface_length = len(fully_adsorption_surface_profile)
	second_layer_list = []
	first_layer_list = []
	for i in range(surface_length):
		site_record = fully_adsorption_surface_profile[i]
		if len(site_record)==1:
			second_layer_list.append('')
			first_layer_list.append(site_record[0])
		else:
			second_layer_list.append(site_record[1])
			first_layer_list.append(site_record[0])
	print(*second_layer_list, sep='\t')
	print(*first_layer_list, sep='\t')

	return()


def simulate_one_heat_cycle(molecule_list, surface_length, volume, D_site_prob, mismatch_prob, kinase_dict, ligase_dict, base_collision_number_per_residue_per_cycle, spontaneous_kination_prob, kination_RF_ratio, spontaneous_ligation_prob, ligation_RF_ratio):
	[fully_adsorption_surface_profile, unused_molecule_count_dict_with_mismatchable_residue] = Stochastic_cover_surface_with_molecules_until_saturation(molecule_list, surface_length, mismatch_prob, D_site_prob)

	#print(fully_adsorption_surface_profile)

	unused_molecule_count_dict_normal_name = remove_MN_labels_from_free_molecules(unused_molecule_count_dict_with_mismatchable_residue)
	fully_adsorption_surface_profile = remove_MN_labels_from_surface_profile(fully_adsorption_surface_profile)

#==============================================================
#	print('#=========================================================================================*')
#	formatted_print_two_layer_profile(fully_adsorption_surface_profile)
#	print('#=========================================================================================*')
#==============================================================

	[kination_deactivation_border_list, ligation_breakage_border_list] = scan_borders_and_get_reaction_types(fully_adsorption_surface_profile)
	[kinase_collision_number_dict, ligase_collision_number_dict, total_kinase_collision_number, total_ligase_collision_number, total_natural_kination_catalyst_collision_number, total_natural_ligation_catalyst_collision_number] = calculate_numbers_of_collisions(base_collision_number_per_residue_per_cycle, unused_molecule_count_dict_normal_name, kinase_dict, ligase_dict, volume, natural_kination_catalyst_collisions_per_volume_per_cycle, natural_ligation_catalyst_collisions_per_volume_per_cycle)
	fully_adsorption_surface_profile = semi_Gillespie_determine_final_status(surface_length, fully_adsorption_surface_profile, kination_deactivation_border_list, ligation_breakage_border_list, kinase_collision_number_dict, ligase_collision_number_dict, total_kinase_collision_number, total_ligase_collision_number, kinase_dict, ligase_dict, spontaneous_kination_prob, kination_RF_ratio, spontaneous_ligation_prob, ligation_RF_ratio, total_natural_kination_catalyst_collision_number, total_natural_ligation_catalyst_collision_number)

#==============================================================
#	formatted_print_two_layer_profile(fully_adsorption_surface_profile)
#	print('#=========================================================================================*')
#==============================================================

	free_molecules_list = release_molecules_from_surface(fully_adsorption_surface_profile, unused_molecule_count_dict_normal_name)

	#print(convert_list_to_count_dict(free_molecules_list))

	return(free_molecules_list)


def simulate_multiple_heat_cycles(cycle_num, molecule_list, surface_length, volume, D_site_prob, mismatch_prob, kinase_dict, ligase_dict, base_collision_number_per_residue_per_cycle, spontaneous_kination_prob, kination_RF_ratio, spontaneous_ligation_prob, ligation_RF_ratio):#, heat_shock_interval):
	for i in range(cycle_num):
		molecule_list = simulate_one_heat_cycle(molecule_list, surface_length, volume, D_site_prob, mismatch_prob, kinase_dict, ligase_dict, base_collision_number_per_residue_per_cycle, spontaneous_kination_prob, kination_RF_ratio, spontaneous_ligation_prob, ligation_RF_ratio)

		#======================================
		# for one cycle one reactor
		molecule_count_dict = convert_list_to_count_dict(molecule_list)
		sorted_mol_name_list = []
		print_dict = {}
		for mol in molecule_count_dict:
			mol_num = molecule_count_dict[mol]
			printout = str(mol)+':'+str(mol_num)
			print_dict[mol] = printout
			sorted_mol_name_list.append(mol)
		
		sorted_mol_name_list.sort()
		sorted_mol_name_list = sorted(sorted_mol_name_list, key=len)
		print_list = []
		for mol in sorted_mol_name_list:
			printout = print_dict[mol]
			print_list.append(printout)
		#print_list.sort()
		print(*print_list, sep=', ')
		#=======================================

	return(molecule_list)


initial_conc_file_name = sys.argv[1]
reactor_index = int(sys.argv[2])# starting from 0

initial_conc_file = open(initial_conc_file_name,'r')
string_record = ''
line_index = 0

for line in initial_conc_file:
	line=line.strip()
	if line_index==reactor_index:
		string_record = line
		break
	else:
		line_index = line_index+1
initial_conc_file.close()


#test_initial_mol_dict = {}

molecule_list = []

count_list = string_record.split(', ')
for name_count in count_list:
	[name,count] = name_count.split(':')
	count = int(count)
	#test_initial_mol_dict[name] = count
	for i in range(count):
		molecule_list.append(name)


molecule_list = simulate_multiple_heat_cycles(cycle_num, molecule_list, surface_length, volume, D_site_prob, mismatch_prob, kinase_dict, ligase_dict, base_collision_number_per_residue_per_cycle, spontaneous_kination_prob, kination_RF_ratio, spontaneous_ligation_prob, ligation_RF_ratio)

#[fully_adsorption_surface_profile, unused_molecule_count_dict_with_mismatchable_residue] = Stochastic_cover_surface_with_molecules_until_saturation(test_molecule_name_list, surface_length, mismatch_prob, D_site_prob)


#print(fully_adsorption_surface_profile)
#print(unused_molecule_count_dict_with_mismatchable_residue)
```
