## Supplemental Python Scripts for "Conditions enabling the persistence of cooperating ribozymes without cellular encapsulation": UpdateGen.py

```
text_file_names = ["script_list.txt", "MergeDistri_list.txt"]


for f in text_file_names:
	parameter_list = []
	file = open(f, "r")
	contents = file.read()
	parameter_list = contents.split(",")
	#print(parameter_list)
	current_generation = parameter_list[0]
	updated_gen_num = int(current_generation) + 1
	parameter_list[0] = str(updated_gen_num)
	print_line = ",".join(parameter_list)
	file.close()
	file = open(f, "w")
	file.write(print_line)
	file.close()
```
