## Supplemental Python Scripts for "Conditions enabling the persistence of cooperating ribozymes without cellular encapsulation": UpdateGen.sh

```
#!/bin/bash

# untar your Python installation. Make sure you are using the right version!
# (optional) if you have a set of packages (created in Part 1), untar them also

# make sure the script will use your Python installation, 
# and the working directory as its home location
export PATH=$PWD/python/bin:$PATH
export PYTHONPATH=$PWD/packages
export HOME=$PWD

# run your script
python3 UpdateGen.py

#exit 1
```
